# Kinome-wide RNAi screen uncovers role of Ballchen in maintenance of gene activation by trithorax group in *Drosophila*

**DOI:** 10.1101/2020.11.26.399824

**Authors:** Muhammad Haider Farooq Khan, Jawad Akhtar, Zain Umer, Najma Shaheen, Ammad Shaukat, Aziz Mithani, Saima Anwar, Muhammad Tariq

## Abstract

Polycomb group (PcG) and trithorax group (trxG) proteins are evolutionary conserved factors that contribute to cell fate determination and maintenance of cellular identities during development of multicellular organisms. The PcG behaves as repressors to maintain heritable patterns of gene silencing and trxG act as anti-silencing factors by maintaining activation of cell type specific genes. Genetic and molecular analysis has revealed extensive details about how different PcG and trxG complexes antagonize each other to maintain cell fates, however the cellular signaling components that contribute to maintenance of gene expression by PcG/trxG remain elusive. Here, we report an ex vivo kinome-wide RNAi screen in *Drosophila* aimed to identify cell signaling genes that facilitate trxG to counteract PcG mediated repression. From the list of trxG candidates, Ballchen (BALL), a histone kinase, known to phosphorylate histone H2A at threonine 119 (H2AT119p), was characterized as a trxG regulator. The *ball* mutant exhibit strong genetic interaction with *Polycomb* (*Pc*) and *trithorax* (*trx*) mutants and loss of BALL also affects expressions of trxG target genes in *ball* mutant embryos. BALL co-localizes with Trithorax on chromatin and depletion of BALL results in increased H2AK118 ubiquitination, a histone mark central to PcG mediated gene silencing. Moreover, analysis of genome-wide binding profile of BALL shows an overlap with 85% known binding sites of TRX across the genome. Both BALL and TRX are highly enriched at actively transcribed genes, which also correlate with presence of H3K4me3 and H3K27ac. We propose that BALL mediated signal positively contributes to the maintenance of gene activation by trxG by counteracting the repressive effect of PcG.

## Introduction

In metazoans, specialization of cell types that make up an organism is linked to cell type specific gene expression patterns established during early development. In order to maintain a specialized state, the particular expression profiles of genes need to be transmitted to daughter cells through successive mitotic divisions in all cell lineages, the phenomenon termed as transcriptional cellular memory. Maintenance of transcriptional cellular memory and consequent cellular identity involves a combinatorial act of various epigenetic mechanisms, such as DNA methylation (Holliday and Pugh, 1975), histone modifications (Allfrey et al., 1964), non-coding RNAs (Jones et al., 1999; Matzke et al., 2001; Volpe et al., 2002) and chromatin remodeling (Sudarsanam and Winston, 2000; Narlikar et al., 2002). Genetic analysis in *Drosophila* uncovered two groups of evolutionarily conserved genes, namely Polycomb group (PcG) and trithorax group (trxG), responsible for maintaining stable and heritable states of gene repression and activation, respectively (Lewis, 1978; Capdevila and García-Bellido, 1981). Molecular analysis revealed that proteins encoded by the PcG and trxG act in large multi-protein complexes, and modify the local properties of chromatin to maintain expression patterns of their target genes. Both groups exert their functions by binding to particular chromosomal elements known as PREs (Polycomb Response Elements) and by interacting with the histones and transcription machinery (Kassis et al., 2017; Cavalli and Heard, 2019). The PcG complexes, PRC1 and PRC2 (Polycomb Repressive Complex 1 and 2), are known to maintain repression by generating ubiquitination of histone H2A at lysine 118 (H2AK118ub1) (Wang et al., 2004) and methylation of histone H3 at lysine 27 (H3K27me) (Francis et al., 2001; Cao et al., 2002; Czermin et al., 2002), respectively. In contrast to PcG, trxG is quite heterogeneous and comprises of proteins that activate transcription by modifying histone tails and ATP dependent chromatin remodeling (Piunti and Shilatifard, 2016). One cellular function that unifies this diverse group of proteins is their role in counteracting PcG mediated gene silencing.

The fact that trxG and PcG coexist at the chromatin regardless of gene expression states of their target genes suggests that PcG and trxG not only compete but also associate with their target genes as dynamic complexes (Breiling et al., 2001; Dellino et al., 2004; Klymenko and Müller, 2004; Papp and Müller, 2006; Beisel et al., 2007). Although, the chromatin structure and modifications appear to play a fundamental role in the maintenance of transcriptional cellular memory, the signal that favors PcG or trxG to either repress or activate remains elusive. It is plausible to assume that cell signaling pathways can be of prime importance due to their ability to respond to intra and extracellular changes as well as their capacity to influence nuclear factors involved in gene repression or activation. Cell signaling components, especially the protein kinases, control a repertoire of cellular processes by modifying more than two-third of cellular proteins function (Ardito et al., 2017), but both the PcG and trxG complexes lack kinases. In *Drosophila,* FSH is the only kinase present in canonical trxG members. Being an atypical kinase with no known kinase domain (Chang et al., 2007), FSH too performs its known cellular functions via its bromodomain and interaction with ASH1 (Kockmann et al., 2013). Although different cellular processes linked to epigenetic inheritance, such as maintenance of chromosomal architecture to transcription (Stadhouders et al., 2019) are regulated by protein kinases (Nowak and Corces, 2000, 2004), the role of cell signaling components in maintaining gene activation by trxG or repression by PcG remains elusive.

Here, we report an RNA interference (RNAi) based reverse genetics screen to identify cell signaling proteins that contribute to the maintenance of gene activation by trxG. An ex vivo kinome-wide RNAi screen was carried out using a well-characterized reporter in *Drosophila* cells (Umer et al., 2019). The primary RNAi screen led to the identification of 27 cell signaling genes that impaired expression of reporter similar to *trx* and *ash1*, two known trxG genes. The majority of the candidates were protein kinases, but regulatory subunits of kinase complexes, kinase inhibitors, nucleotide kinases and a few lipid kinases, were also present in the list. Remarkably, presence of FSH, the only trxG member with predicted kinase activity, in the list of candidates validated functionality of the screen. From the list of candidates, selected serine-threonine kinases were further confirmed in a secondary screen which affected reporter system similar to the effect of TRX and ASH1 depletion.

Next, we performed genetic and molecular analysis of Ballchen (BALL), a histone kinase in the list of candidate genes, and showed that BALL is required to maintain gene activation by trxG. BALL mutant exhibit trxG like behavior by strongly suppressing extra sex comb phenotype caused by *Pc* mutations as well as by enhancing the homeotic phenotype in *trx* mutants. This strong genetic interaction between *ball* and trxG system corroborates with a drastic reduction in expression of homeotic and non-homeotic targets of trxG in *ball* mutant embryos. Importantly, reduced expression of trxG targets due to depletion of BALL also correlates with enhanced levels of H2AK118ub1, a histone modification central to PcG mediated gene repression (Blackledge et al., 2014, 2020; Kalb et al., 2014; Tamburri et al., 2020). Finally, the genome-wide binding sites of BALL identified in *Drosophila* S2 cells by ChIP-seq revealed BALL occupancy at transcription start sites and it binds at 85% of known TRX binding sites. Since BALL is known to phosphorylate H2A threonine 119 (H2AT119p) (Aihara et al., 2004), our data supports the notion that BALL contributes to maintenance of gene activation by counteracting PRC1 mediated H2AK118ub1.

## Results

### Kinome-wide RNAi screen discovered kinases affecting cell memory maintenance

To identify the role of cell signaling genes in the maintenance of gene activation by trxG, we used a previously characterized cell-based reporter *PBX-bxd-IDE-F.Luc* (hereafter referred to as *PRE-F.Luc*). In this reporter, the *Firefly* luciferase gene is under the control of *Drosophila Ubx* (*Ultrabithorax*) promoter and *bxd PRE* along with *PBX* (*postbithorax*) and *IDE* (*Imaginal Disc Enhancer*) enhancers of *Ubx*. The sensitivity and specificity of this reporter to the changing levels of PcG and trxG are already described (Umer et al., 2019). Using this reporter, we performed an ex vivo RNAi screen, covering all known and predicted kinases and their associated proteins from the HD2 dsRNA library (Horn et al., 2010). Each gene was knocked down in triplicates and the entire experiment was performed twice. dsRNAs against known trxG members (*trx, ash1*) and the reporter gene (*F.Luc*) were used as positive controls, whereas dsRNA against *LacZ* or *GFP* was used as a negative control in all plates. *Drosophila* cells treated with dsRNAs were transfected with *PRE-F.Luc* reporter along with actin promoter-driven *Renilla* luciferase (*R.Luc*), used as a normalization control (Figure 1A, S1). Since *R.Luc* is driven by a constitutive promoter, it also served as a control to exclude genes that may affect general transcription of genes. After five days of transfection, the activity of both *F.Luc* and *R.Luc* was determined and Z-scores were calculated. Based on the Z-scores obtained from positive controls (*trx, ash1*), cut-offs were defined and a list of potential trxG regulators was generated (Table 1). Of 400 genes that were screened, 27 specifically resulted in reduced *PRE-F.Luc* expression, an effect similar to the knockdown of *trx* and *ash1*. The only trxG member with predicted kinase activity, *fs(1)h* was also found among top hits of the screen. Moreover, proteins having known genetic and molecular interactions with trxG also appeared in the list of candidates. For instance, human CDK1, a candidate gene in our list, is known to phosphorylate EZH2, enzymatic subunit of PRC2, at threonine 487 and as a consequence decreases its methyltransferase activity in mesenchymal stem cells (Wei et al., 2011). In addition to protein kinases, lipid kinases, nucleotide kinases, kinase inhibitors and regulatory subunits of kinase complexes were also present in the list of candidates. Gene ontological analysis using STRING database revealed that most candidates reside in nucleus and many interact with each other in cell cycle regulation (Figure 1B). For example, CDK2 along with associated Cyclin E are in cell cycle regulators cluster that also physically interact with trxG proteins, Brahma (BRM) and Moira (MOR), the core proteins of Brahma associated protein complex (Brumby et al., 2002).

**Figure 1.**
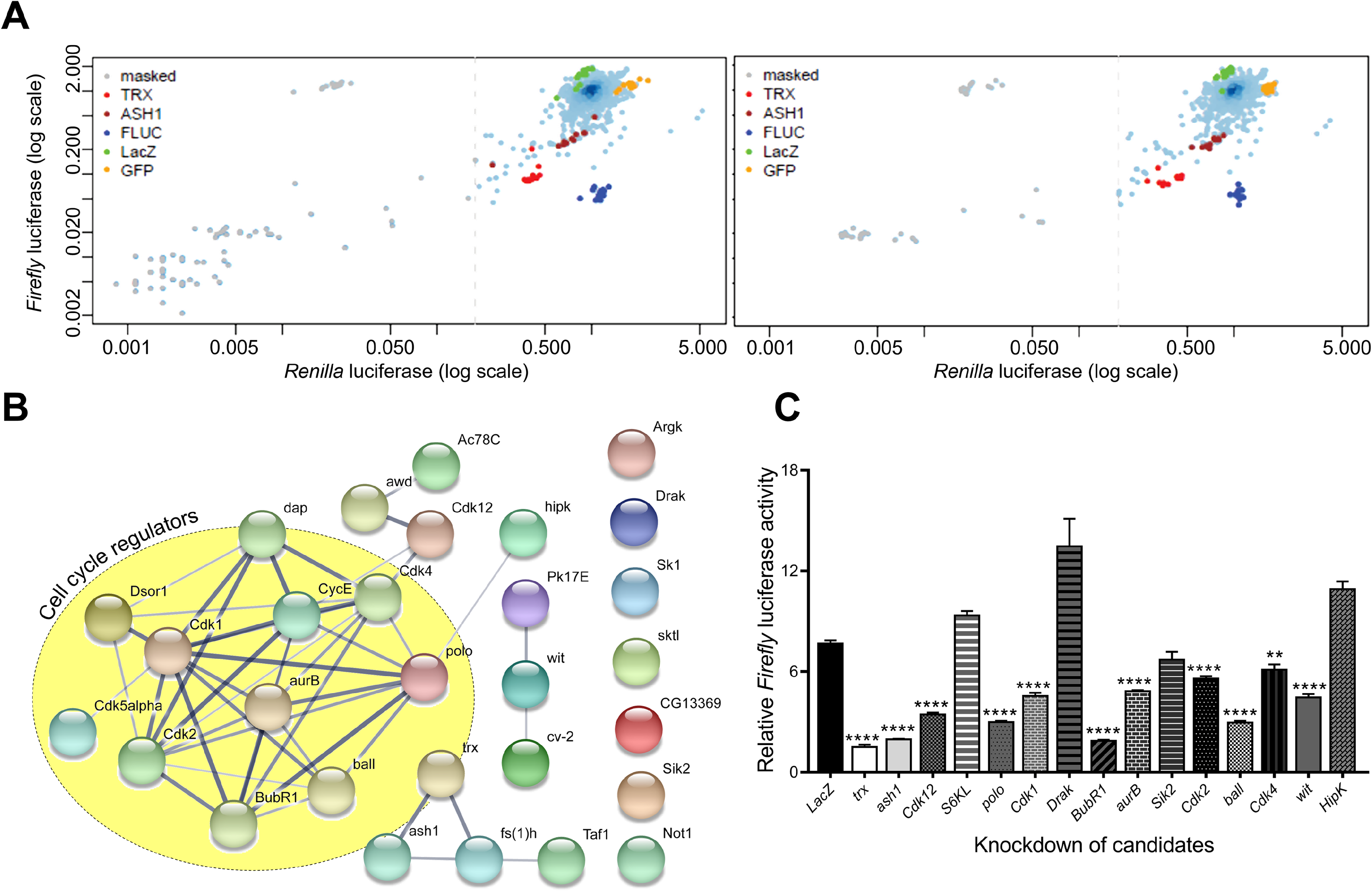
Kinome-wide RNAi screen data analysis and validation. (A) Scatterplots of the plate median corrected intensity values for *Firefly* luciferase (F.Luc) against the plate median corrected intensity values for *Renilla* luciferase (R.Luc). To generate a list of trxG candidate genes that affect gene activation, Z-scores of *trx* (red) and *ash1* (maroon) knockdown were used to set cut-off value which is represented by a dashed line. Knockdown of *trx* (red) and *ash1* (maroon) as well as knockdown of *F.Luc* (blue) were used as positive controls. Cells treated with dsRNA against *LacZ* (green) or *GFP* (orange) served as negative controls. Genes affecting activity of both F.Luc and R.Luc were masked (gray) and were not investigated further. Data shown represents two independent experiments of the kinome-wide RNAi screen. (B) Protein-protein interaction network of candidate genes, generated using STRING database (Snel, 2000; Szklarczyk et al., 2019). Nodes depict proteins that are connected by lines of varying thickness. Thickness of lines demonstrates the degree of confidence for the interaction between connected nodes. The yellow circle is marking candidates that are known to associate with cell cycle regulation. (C) Knockdown of selected serine-threonine kinases from list of candidate genes from primary screen resulted in significantly lower relative *Firefly* luciferase activity when compared with cells treated with dsRNA against *LacZ*. Cells treated with dsRNA against *trx* and *ash1* were used as positive control. Experiment was performed in triplicate and independent t-tests were done for analysis (** p ≤ 0.01 or **** p≤ 0.0001).

**Table 1.**
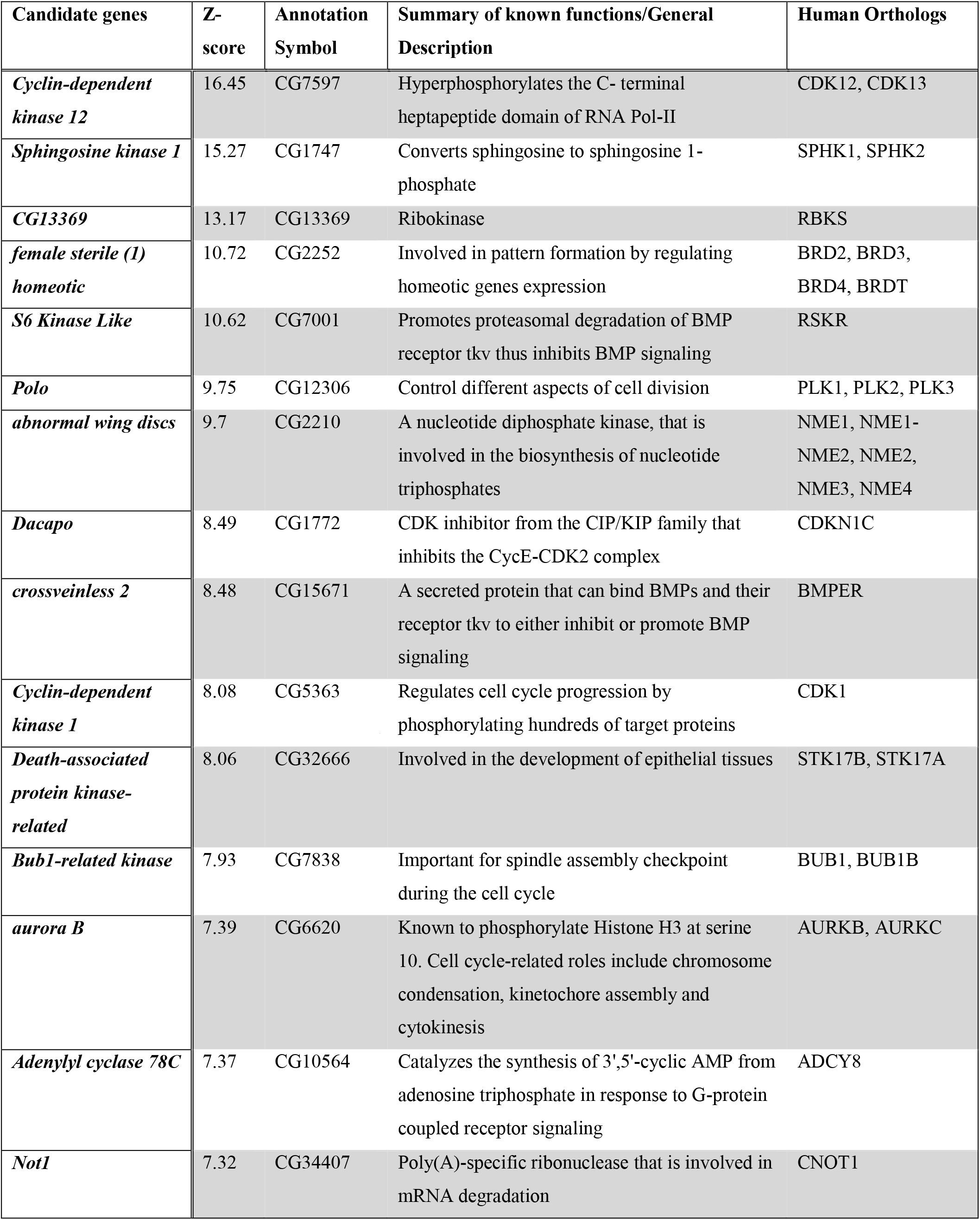
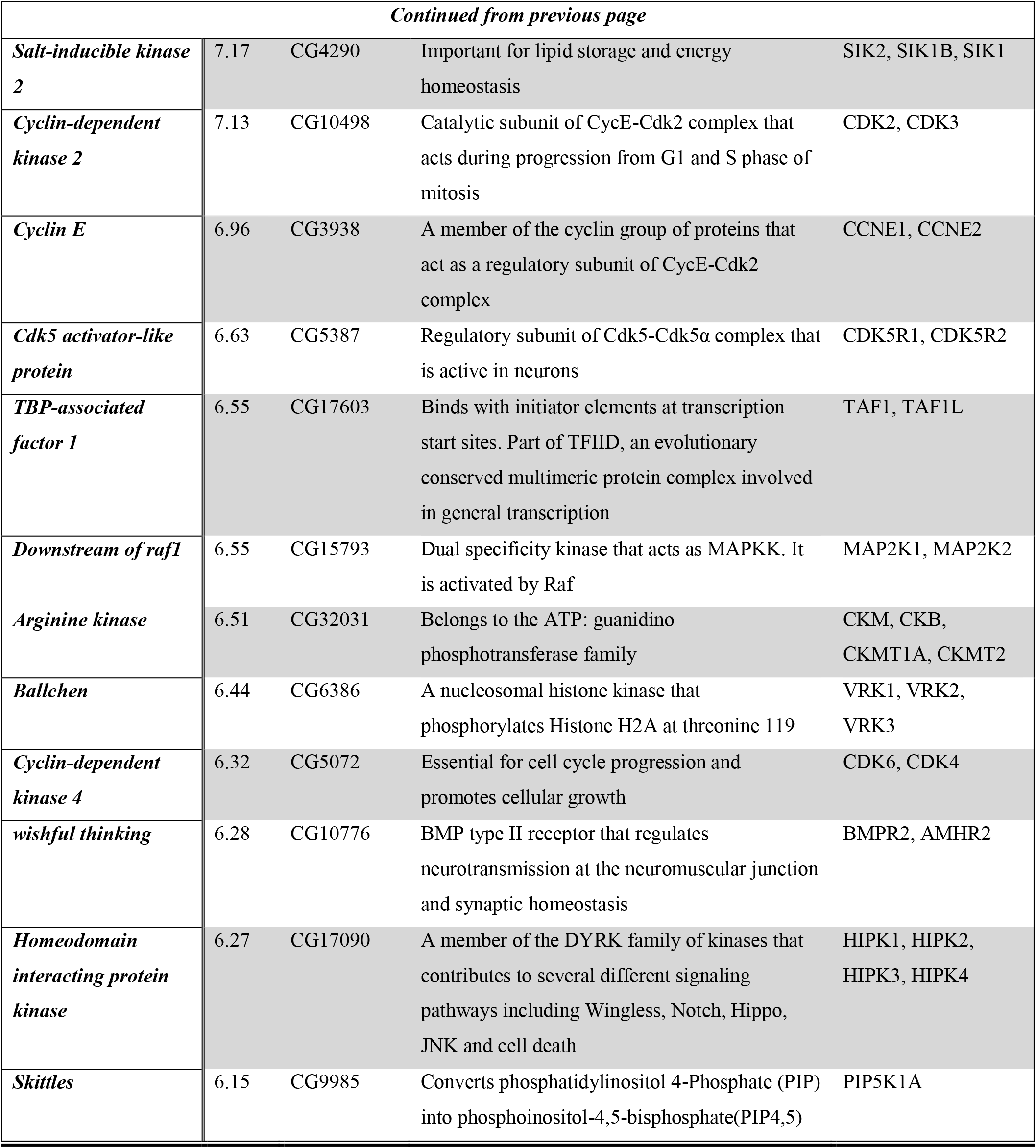
List of candidate genes along with their respective Z-scores, annotation symbols, human orthologs and a summary of their known function. General description of each gene is based on information obtained from Uniprot and Flybase databases.

Next, we selected serine/threonine protein kinases from the list of candidates and performed a secondary screen using *PRE-F.Luc* (Figure 1C). Cells treated with dsRNA against *trx* and *ash1* were used as positive controls while cells treated with dsRNA against *LacZ* served as a negative control. Depletion of nine of thirteen selected kinases in secondary screen resulted in significantly decreased activity of *F.Luc*. Since dsRNA amplicons used to knockdown target genes in the secondary screen were different from the ones used in the primary screen, it further validated our results from the primary screen. Finally, we chose *ballchen* (*ball*) from the list of candidate genes to explore its genetic and molecular link with trxG because it is a known histone kinase which phosphorylates histone H2AT119 (Aihara et al., 2004).

Depletion of *ball* drastically reduced expression of the reporter gene, relative F.Luc activity, when compared with other candidates in the secondary screen. Moreover, it is also known to be involved in both cell cycle (Fiona Cullen et al., 2005) and signal transduction pathways (Herzig et al., 2014; Yakulov et al., 2014), making it the most suitable representative gene of the candidates list.

### *ball* exhibits *trxG* like behavior

To investigate whether *ball* genetically interacts with the PcG/trxG system, *ball* mutant flies were crossed with two different mutant alleles of *Pc* (*Pc^1^*, *Pc^XL5^*). *Pc* heterozygous mutants display a strong extra sex comb phenotype in males. The *ball* mutant (*ball^2^*) strongly suppressed this extra sex comb phenotype (Figure 2A, B) which supports the role of *ball* as a trxG-like factor controlling homeotic phenotype. Next, we examined the genetic interaction of *ball* with *trithorax (trx)* by crossing *ball^2^* with two different alleles of *trx* (*trx^1^* or *trx^E2^*) mutants. As compared to wild-type flies, *trx* heterozygous males show loss of pigmentation on 5^th^ abdominal segment referred to as A5 to A4 transformation (Figure 2C) and *ball^2^/trx* trans-heterozygotes exhibit significantly more A5 to A4 transformations (Figure 2C) as compared to *trx* heterozygotes indicating strong enhancement of *trx* phenotype by *ball*. The suppression of extra sex combs phenotype and enhancement of *trx* phenotype by *ball* mutation indicate that *ball* exhibits a trxG like behavior.

**Figure 2.**
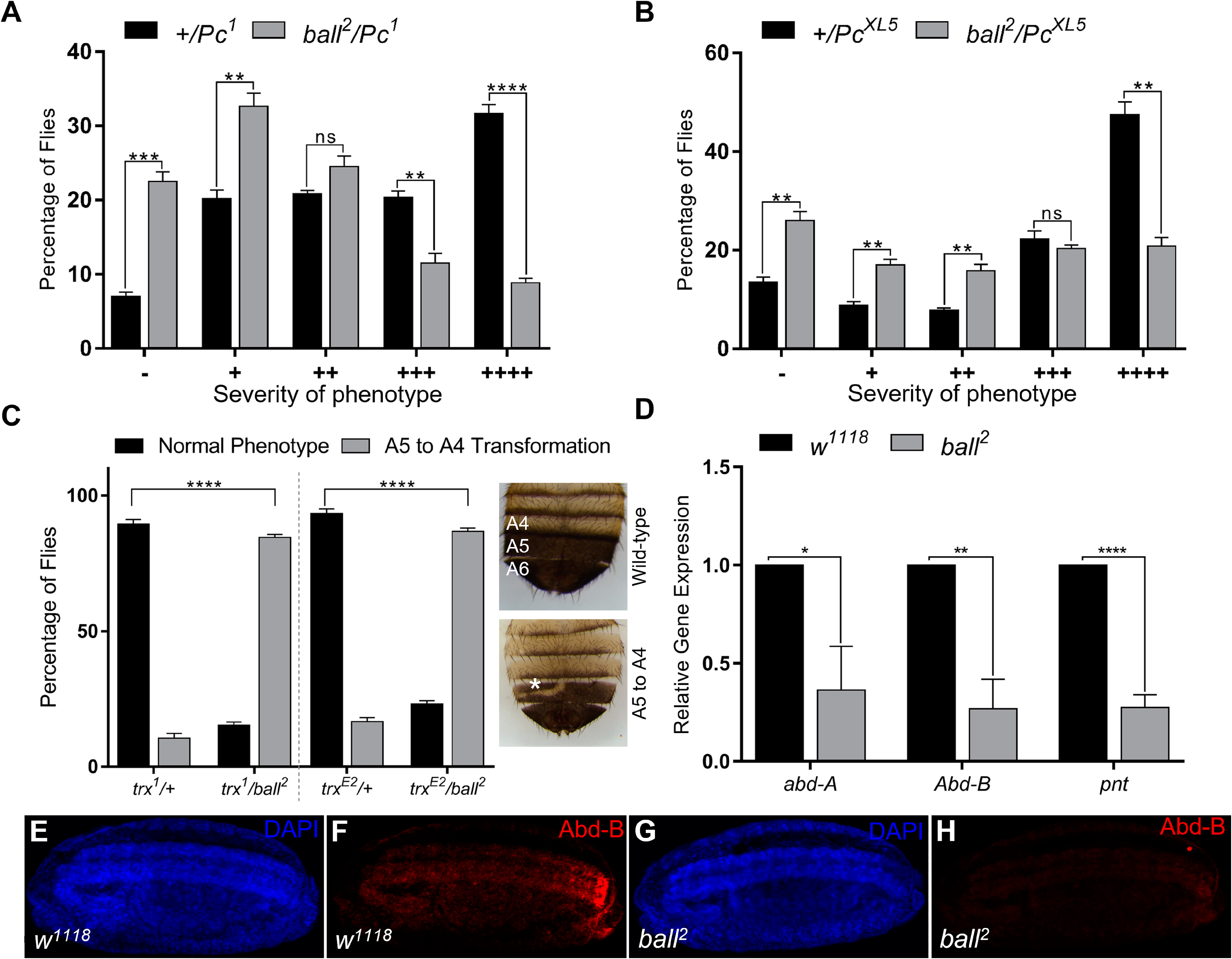
*ballchen* mutation exhibits trxG like behavior. (A, B) Ballchen mutant flies (*ball^2^*) were crossed with two different alleles of *Pc* (*Pc^1^* and *Pc^XL5^*), while *Pc* alleles (*Pc^1^* and *Pc^XL5^*) crossed with *w^1118^* flies were used as control. Heterozygous *Pc/+* males, *Pc^1^*/+ (A) and *Pc^XL5^*/+ (B), from control crosses exhibit strong extra sex combs phenotype. In contrast, *ball^2^* strongly suppressed both the *Pc* alleles to none or few extra sex combs in *ball^2^/Pc^1^* (A) as well as *ball^2^/Pc^XL5^* (B) flies. 200 male flies were analyzed for each cross and data shown represents two independent experiments. Based on strength of phenotype, male flies were categorized according to the severity of extra sex combs phenotype. These categories are: -, no extra sex combs; +, 1-2 hairs on 2^nd^ leg; ++, more than 3 hairs on 2^nd^ leg; +++, more than 3 hairs on 2^nd^ leg and 1-2 hairs on 3^rd^ leg; ++++, strong sex combs on both 2^nd^ and 3^rd^ pairs of legs. (C) *ball^2^* mutant flies were crossed to two different alleles of *trx* (*trx^1^* and *trx^E2^*), and *ball^2^/trx* males in F1 were scored for loss of pigmentation in A5 abdominal segment (A5 to A4 transformation, marked with asterisk). The *trx* alleles (*trx^1^* and *trx^E2^*) crossed with *w^1118^* flies i.e. *trx^1^/+* and *trx^E2^/+*, show A5 to A4 transformation when compared to wild type flies. Strong enhancement of A5 to A4 transformation can be seen in *ball^2^/trx* double mutants when compared to *trx/+*. Representative images for wild-type phenotype (top right) and A5 to A4 transformation (marked by white asterisk) are presented for *trx/ball* male flies (bottom right). All crosses were done in triplicates and independent t-tests were performed for analyzing each category (* p ≤ 0.05, ** p ≤ 0.01, *** p≤ 0.001 or **** p≤ 0.0001). (D) Significantly low levels of *abd-A*, *Abd-B* and *pnt* expression was detected through qRT-PCR in homozygous *ball^2^* embryos when compared with *w^1118^* embryos. Independent t-tests were performed for each gene analysis (* p ≤ 0.05, ** p ≤ 0.01, *** p≤ 0.001 or **** p≤ 0.0001). (E-H) Immunostaining of stage 15 embryos with anti-Abd-B is shown in *w^1118^* (E, F) as well as homozygous *ball^2^* (G, H) embryos. As compared to *w^1118^* (F), *ball^2^* embryos showed strongly diminished Abd-B (H) expression.

Next, we investigated the effect of *ball* mutation on expression of homeotic and non-homeotic targets of PcG/trxG. Since homozygous *ball* mutants do not develop into adults (Herzig et al., 2014), we used homozygous *ball* embryos to analyze mRNA levels of *abd-A, Abd-B,* and *pnt* through real-time PCR (Figure 2D). As compared to *w^1118^* embryos, a significant reduction in expression of *abd-A, Abd-B* and *pnt* was observed. Since *Abd-B* is the gene responsible for pigmentation in A5 and A6 abdominal segments in males (Jeong et al., 2006), its depletion in *ball* mutant embryos corroborates with loss of pigmentation in *ball^2^/trx* trans-heterozygotes. It was further validated by immunostaining of stage 15 homozygous *ball^2^* embryos with anti-Abd-B antibody (Figure 2E-H). In wild-type embryos at this stage of development, Abd-B expression progressively increases from PS10-14 (Figure 2F). However, in *ball^2^* mutants, Abd-B expression is drastically reduced (Figure 2H) which correlates with significantly diminished *Abd-B* mRNA levels in homozygous *ball^2^* embryos (Figure 2D). Together with genetic evidence, these results suggest that trxG require BALL for maintenance of gene activation during development.

### BALL co-localizes with TRX and inhibits PRC1

BALL is known to bind chromatin during mitosis, however its association with chromatin during interphase is not yet established. We show that immunostaining of polytene chromosomes from third instar larvae with anti-BALL antibody revealed association of BALL on a number of sites on chromosomes (Figure 3A-C). Next, to investigate if BALL and TRX co-localize on chromatin, *UAS-ball-EGFP* transgenic flies were crossed to a salivary gland specific GAL4 driver line (*sgs-gal4*). Immunostaining of polytene chromosomes from salivary glands of third instar larvae expressing *ball-EGFP* revealed a substantial but partial overlap between BALL and TRX (Figure 3D-F). Besides co-localizing with TRX, BALL was seen to associate with couple of hundred sites across the polytene chromosomes (Figure 3B, D). Chromatin association of BALL at TRX binding sites was validated by ChIP analysis from stable cells expressing the *ball* coding sequence fused with FLAG epitope under copper inducible promoter (Figure 3G). BALL was found at the transcription start sites (TSS) of *pipsqueak* (*psq*), *pannier* (*pnr*), *pointed* (*pnt*) and *disconnected* (*disco*) which are known binding sites of PcG/trxG. Moreover, an association of BALL was also observed at *iab-7PRE*, *bxd* and *Dfd* regulatory regions of homeotic genes (Figure 3G). In contrast, ChIP from empty vector control cells using anti-FLAG antibody resulted in negligible enrichment at all the regions analyzed. Additionally, chromatin association of BALL on TRX binding sites was also analyzed with antibody against endogenous BALL using wild-type *Drosophila* S2 cells. A stronger enrichment of BALL, but with a similar pattern, was seen on all the non-homeotic and homeotic targets analyzed (Figure 3G).

**Figure 3.**
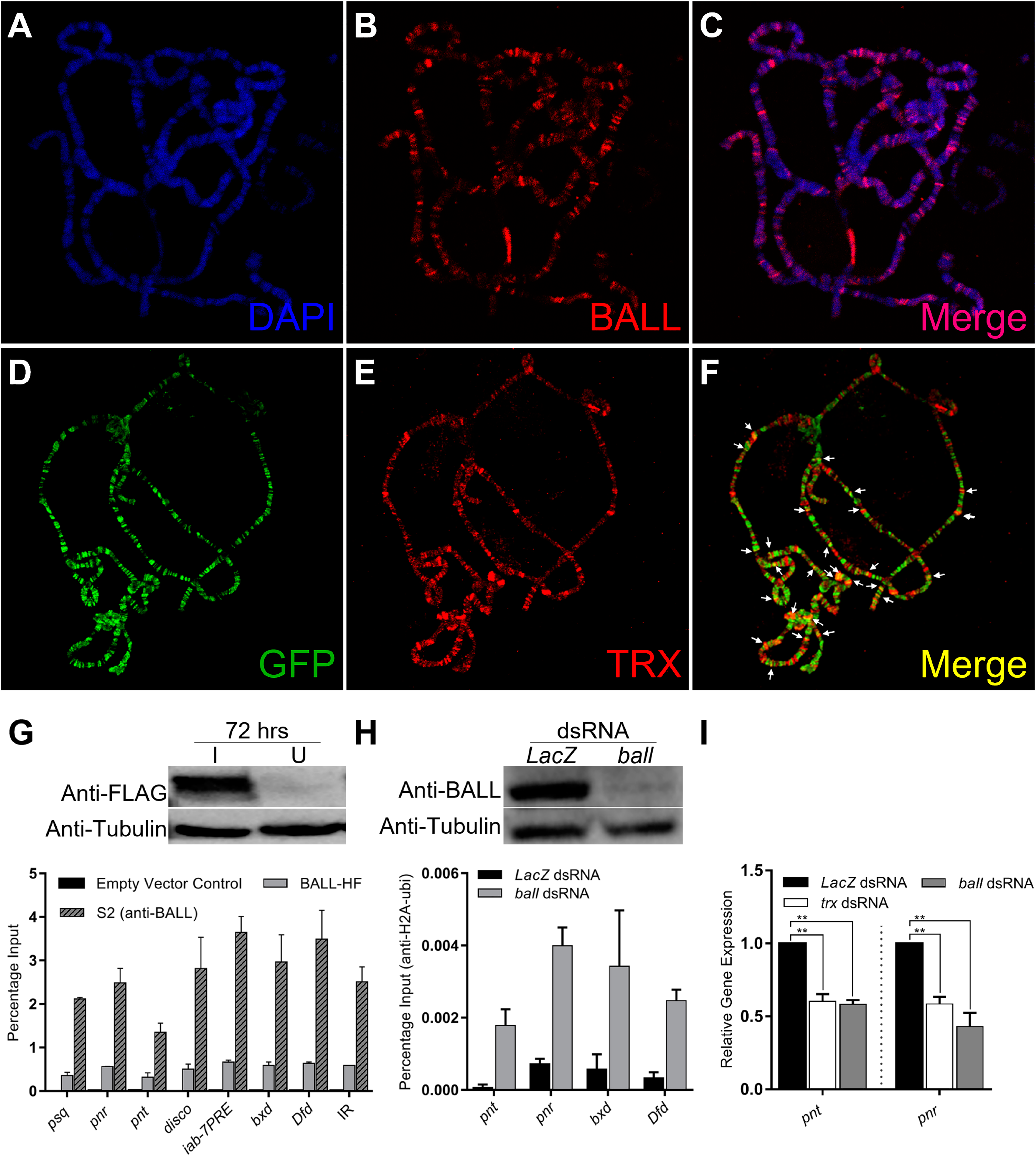
BALL co-localizes with TRX at the chromatin and antagonizes PcG. (A-C) Polytene chromosomes from third instar larvae stained with DAPI (A) and anti-BALL (B) antibody. (D-F) Third instar larvae expressing GFP-tagged BALL were stained with anti-GFP (D) and anti-TRX (E) antibodies. Co-localization of BALL and TRX is clearly seen at several loci (F, white arrows). (G, top panel) Western blot analysis of *Drosophila* S2 stable cells, expressing FLAG-tagged BALL (*ball-HF*) protein only in the induced sample (I) as compared to un-induced (U) cells. (G, bottom panel) Strong enrichment of BALL at trxG targets was observed by ChIP using anti-FLAG antibody in *Drosophila* S2 cells expressing FLAG-tagged BALL (BALL-HF). ChIP from empty-vector control cell line using anti-FLAG antibody was used as control. ChIP from wild-type S2 cells using anti-BALL antibody also showed similar but stronger enrichment. (H, top panel) Knockdown of *ball* drastically decreased the amount of BALL protein expressed when compared to cells treated with *LacZ* dsRNA. (H, bottom panel) ChIP from BALL depleted cells showed enhanced H2AK118ub1 levels at non-homeotic (*pnt*, *pnr*) and homeotic (*bxd*, *Dfd*) targets of trxG, as compared to cells treated with dsRNA against *LacZ*.

Since BALL is known to phosphorylate histone H2AT119, a residue adjacent to H2AK118, we questioned whether the H2AT119ph mark is inhibitory for H2AK118ub1, an established hallmark of PcG mediated gene repression (Blackledge et al., 2014, 2020; Kalb et al., 2014; Tamburri et al., 2020). To this end, we performed ChIP from cells where *ball* was depleted using dsRNA against *ball*. The BALL depleted cells exhibited an increased H2AK118ub1 at both transcriptionally active (*pnt*, *pnr*) and repressed (*bxd*, *Dfd*) targets of trxG as compared to cells treated with dsRNA against *LacZ* which served as a control (Figure 3H). Depletion of *ball* and the efficiency of knockdown was confirmed through Western blotting using anti-BALL antibody (Figure 3H). As the increase in H2AK118ub1 should also reflect on expression levels of the target genes, we analyzed transcriptionally active genes (*pnt* and *pnr*) mRNA levels through qRT-PCR in BALL depleted cells. The depletion of BALL indeed resulted in decreased expression of *pnt* and *pnr* compared to cells treated with dsRNA against *LacZ* (Figure 3I). These results suggest that BALL is required by trxG to maintain gene activation and it may counteract PcG by inhibiting H2AK118ub1, a histone modification catalyzed by PRC1 subunit dRING, and in turn shift the balance in favor of trxG.

### Genome-wide binding profile of BALL correlates with TRX and gene activation

The association of BALL with chromatin and its co-localization with TRX on polytene chromosomes intrigued us to determine genome-wide binding sites of BALL. For this purpose, ChIP with antibody against endogenous BALL was performed using formaldehyde fixed chromatin from *Drosophila* S2 cells and purified DNA was sequenced using BGISEQ-500. BALL showed very similar binding profile to that of TRX as it was found to associate with 85% known binding sites of TRX across the genome (Figure 4A), which further signifies the involvement of BALL in transcriptional cellular memory governed by PcG/trxG system. Further analysis of ChIP-seq data revealed that BALL is present within ±1kb of TSSs (Transcription Start Sites) of genes (Figure 4A). BALL binds to total 6195 sites, details of which are given in Figure S2 and Table S1. Besides an overwhelming overlap between BALL and TRX binding sites, BALL association with chromatin also correlates with H3K4me3 and H3K27ac marks linked to gene activation (Figure 4B, C). Since expression of *pnt* and *pnr* genes is reduced after depletion of BALL or TRX, ChIP-seq data was specifically analyzed for association of BALL and TRX across genomic regions of *pnt* and *pnr*. High levels of both the BALL and TRX at the *pnt* and *pnr* genomic regions coincide with high levels of H3K27ac and H3K4me3 and little to no PC is present at these sites (Figure 4B). Additionally, analysis of BALL binding sites across the bithorax complex (BX-C) genomic region revealed that BALL binding profile mimics known TRX binding sites in BX-C. Since genes within BX-C are silent in S2 cells, their repression correlates with high prevalence of of PC enrichment throughout BX-C whereas both BALL and TRX are present on few overlapping binding sites across the BX-C. Importantly, an actively transcribed gene, CG14609, just downstream of *Ubx* in BX-C show highly enriched BALL and TRX association that correlates with both H3K4me3 and H3K27ac. All these results indicate a close association between BALL and TRX which suggests an important role for BALL in maintenance of gene activation governed by trxG.

**Figure 4.**
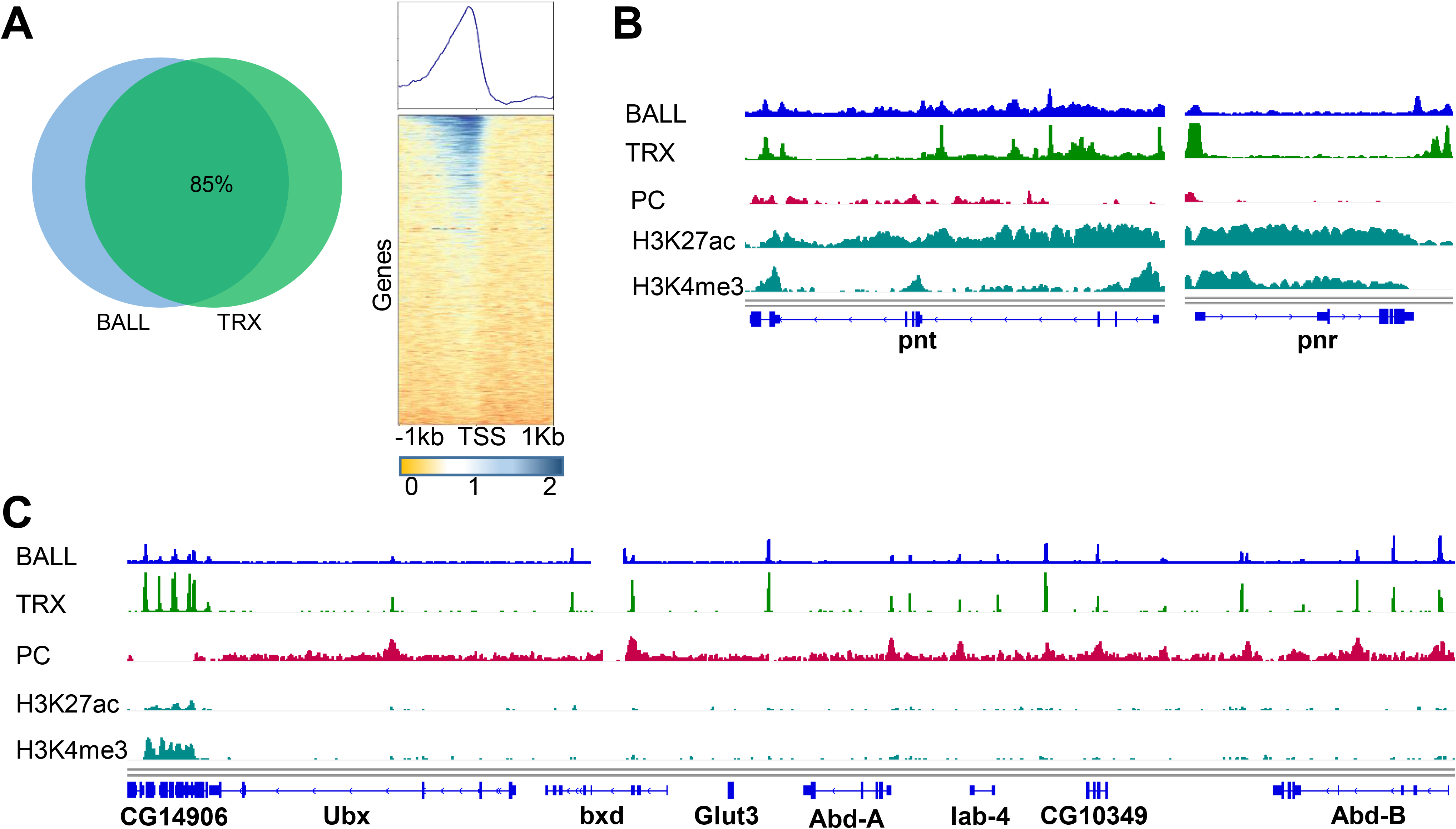
Genome-wide binding profile of BALL analyzed by ChIP-seq from Drosophila S2 cells. (A) Venn diagram analysis showed 85% co-occupancy of BALL on known TRX binding sites. Heat map illustration of BALL binding sites across genome show enrichment of BALL within ± 1kb of TSSs of genes. (B) Genome browser view of BALL association compared with patterns of TRX, PC, H3K27ac and H3K4me3 association at *pnt* and *pnr* genes. (C) Genome browser view of ChIP-seq data showing binding profile of BALL and its comparison with TRX, PC, H3K27ac and H3K4me3 association across the bithorax complex (BX-C). Highly enriched BALL and TRX correlates with high levels of both H3K27ac and H3K4me3 and absence of PC at CG14906 present just downstream of Ubx. In contrast, BX-C genes i.e. *Ubx*, *abd-A* and *Abd-B* which are silent in S2 correlates with highly enriched PC all across BX-C. Both BALL and TRX are present at few overlapping binding sites in BX-C.

## Discussion

Our results demonstrate success of reverse genetics approach to identify cell signaling genes that impact trxG mediated gene activation. With the help of a robust cell based reporter, we identified 27 genes from the kinome-wide RNAi screen in *Drosophila*. This list of candidate genes includes cell signaling kinases and associated genes which were finalized using a stringent criterion based on the Z-scores of *trx* and *ash1* knockdown. Presence of *fs(1)h,* the only trxG member with predicted kinase activity, among the candidate genes validated the robustness of our screen in identifying kinases regulating trxG dependent gene activation. FSH protein facilitates transcription by recognizing acetylated histone marks through its bromodomain and interaction with ASH1 (Kockmann et al., 2013). Other interactors of trxG including *skittles* (*sktl*), *Cyclin E* (*CycE*) and *Cdk2* were also present in the list of candidate genes. SKTL, a nuclear phosphatidylinositol 4-phosphate 5-kinase, is known to interact with trxG member ASH2 (Cheng and Shearn, 2004) and its catalytic product, phosphoinositol-4,5-bisphosphate (PIP4,5), also stabilizes mammalian Brahma associated factors (BAF) complex on the chromatin (Zhao et al., 1998). CycE and its associated kinase CDK2 are also known to interact with BRM and MOR, two known members of trxG, which form core of chromatin remodeling Brahma associated protein (BAP) complex (Brumby et al., 2002).

Protein kinases can also affect epigenetic cellular memory by causing phosphorylation of a repertoire of epigenetic regulators including PcG and trxG proteins. For instance, CDK1, a candidate gene in our list is reported to phosphorylate several proteins involved in histone modifications and transcription regulation. For example, MLL2, LSD1, G9a, SUV39H2, SETD2, DOT1L, p300, KDM2A, HDAC6 are some important histone modifiers included in the list of CDK1 substrates among many others. Additionally, DNA maintenance methyltransferase (DNMT1) as well as Tet1 and Tet2 proteins that reverse DNA methylation are also phosphorylated by CDK1 (Michowski et al., 2020). Cell cycle regulation is a unifying feature of many of our candidates as suggested by the STRING analysis (Figure 1B). Such an involvement of cell cycle regulators in maintenance of epigenetic cellular memory corroborates with the gradual decrease in PRC2 mediated H3K27me3 levels with each round of cell division in flies (Laprell et al., 2017). Similarly differential recruitment of PRC2 subunits to cell lineage-specific promoters is reported across cell cycle in mouse embryonic stem cells (mESCs) (Asenjo et al., 2020). Identification of cell cycle regulators in our screen supports the notion that PcG/trxG system is regulated in a cell cycle dependent manner (Laprell et al., 2017; Asenjo et al., 2020), which may play a role in faithful inheritance of epigenetic states.

Genetic and molecular characterization of BALL, one of the candidate genes from the kinome-wide RNAi screen, described here has revealed novel role for BALL in transcriptional cellular memory. Although, BALL is a known histone kinase that modifies histone H2AT119 (Aihara et al., 2004), it was not associated with the maintenance of gene activation by trxG prior to our kinome-wide RNAi screen. Our results revealed a strong genetic interaction of *ball* mutant with *Pc* and *trx* mutations. We demonstrated that *ball* mutant exhibit trxG like behavior as it strongly suppresses extra sex comb phenotype of *Pc* mutants. Moreover, mutation in *ball* enhances *trx* mutant phenotype, which corroborates with strong suppression of extra sex comb phenotype. This trxG like behavior of *ball* is substantiated by the fact that expression of trxG target genes is reduced in *ball* homozygous mutant embryos. Presence of BALL together with TRX at chromatin suggests that trxG requires BALL for maintenance of gene activation. Since BALL phosphorylates histone H2AT119, a residue adjacent to H2AK118 which is ubiquitinated by dRING in PRC1, it is plausible to assume that BALL mediated phosphorylation may counteract H2AK118ub1 by PRC1 and facilitate trxG. Additionally, enhanced levels of H2AK118ub1 were found at trxG targets when function of BALL was compromised in cells which corroborates with downregulation of trxG target genes. All the results presented here support a notion that BALL mediated phosphorylation of H2AT119 counteracts H2AK118ub1 (Aihara et al., 2004, 2016) and plays a role in gene activation. Interplay between phosphorylation of specific amino acids with covalent modifications of neighboring lysine residues in histones is reported to play a role in epigenetic gene regulation. For example, H3S28 phosphorylation is known to counteract H3K27me3 and promote H3K27ac in maintenance of gene activation by trxG (Gehani et al., 2010). Similarly, H3S10 phosphorylation is proposed to promote H3K14ac and counteract H3K9me3 and contribute to gene activation (Lo et al., 2000; Fischle et al., 2003). Since genome-wide binding profile of BALL also correlates with presence of H3K27ac and H3K4me3 on active genes, it is important to investigate if BALL mediated phosphorylation impacts H3K27ac or H3K4me3. Notably, phosphorylated histone H3 threonine 3 (H3T3p), a mitosis specific mark catalyzed by Haspin kinase (Dai et al., 2005) and mammalian VRK1 (homolog of BALL) (Kang et al., 2007), is reported to inhibit demethylation of H3K4me3 (Su et al., 2016). Interestingly, H3K4me3 is also a known boundary feature of topologically associated domains (TADs) (Dixon et al., 2012), thus such interactions may explain maintenance of active TAD boundaries and epigenetic inheritance across cell division. Moreover, depletion of mammalian VRK1 was shown to result in loss of H3K14 acetylation and H4 acetylation (Salzano et al., 2015). Such a wide-ranging effect of VRK1 on different covalent modifications of histones may well be an indirect consequence of phosphorylation of chromatin binding proteins by VRK1. For example, CREB is also phosphorylated by VRK1 at serine 133 (Kang et al., 2008) and this specific phosphorylation is known to enhance CREB interaction with CBP involved in H3K27ac. This interplay of VRK1, CREB and CBP is known to increase CREB target genes transcription (Radhakrishnan et al., 1998; Dyson and Wright, 2005). Recently, VRK1 is shown to positively regulate DNA damage repair by phosphorylating Tip60 (García-González et al., 2020). However, discovery of mechanistic basis of BALL mediated phosphorylation in maintenance of gene activation by trxG requires further scrutiny at molecular and biochemical levels. Since BALL plays a role in chromosome condensation during mitosis, it will be interesting to probe if phosphorylation mediated signaling events involving BALL may help survival of chromatin structures and modifications involved in the maintenance of transcriptional cellular memory during processes of DNA replication and cell division. Based on the discovery of a predominantly large number of cell cycle associated genes in our screen, it is also interesting to scrutinize if phosphorylation could be the central epigenetic mark that acts as a stable anchor for epigenetic inheritance and help restore dynamic interactions between PcG and trxG proteins and their target genes after cell division.

## Materials and methods

### RNAi screen

HD2 kinome-wide sub-library was used for primary RNAi screen (Horn et al., 2010), details of which can be obtained from http://rnai.dkfz.de. D.Mel-2 cells were incubated with dsRNAs against all known and predicted kinases and associated proteins. In 384 well plates, dsRNAs against each gene was present in triplicates and the entire experiment was performed twice in primary screen. Six 384 well plates were used for each experiment and a final concentration of 50ng/μl dsRNA was used in each well. The *trx, ash1* and *F.Luc* specific dsRNAs were used as positive controls whereas dsRNAs against *LacZ* and *GFP* were used as negative control in each plate. 8000 cells in 30μl were dispensed per well with a multidrop dispenser. The cells were spun down for 10s at 900 rpm and the plates were sealed and incubated at 25°C. Next day, *PRE-F.Luc* and *Actin-R.Luc* were transfected in these cells where *R.Luc* served as a normalization control. After 5 days of transfections, F.Luc and R.Luc values were recorded and ratios of the experimental reporter (F.Luc) to the invariant co-reporter (R.Luc) values were calculated to exclude possible artifacts and rule out genes that may affect general transcription. Knockdown of genes that affected both F.Luc and R.Luc were removed from further analysis. Relative F.Luc expressions were averaged for both replicates. Instead of using 99.99% confidence values as cut-offs, more stringent cut-offs were defined based on the Z-scores of *trx* and *ash1* knockdown. In secondary screen, primers for synthesizing dsRNA were chosen from DRSC library instead of HD2 library that can be found at https://fgr.hms.harvard.edu/fly-cell-based-rnai. Instead of 384 well plates, 96 well plates were used with each well having 2μg dsRNA. 50000 cells in 100μl were dispensed in each well and same experimental procedure described above for primary screen was performed.

### Fly strains and genetic analysis

The following fly strains were obtained from Bloomington *Drosophila* Stock Center: *Pc^XL5^/TM3Ser,Sb, Pc^1^/TM3*Ser, *trx^1^* (BN 2114), *trx^E2^* (BN 24160). *ball^2^* was a gift from A. Herzig (Herzig et al., 2014). The strain used for gene expression analysis in embryos was: *P{neoFRT}82B e ball^2^/TM3, P{w[+mC]=ActGFP}JMR2, Ser[1]* (Referred to as *ball^2^/GFP*). For immunostaining of polytene chromosomes *P{w^+mC^UASp-ball.T:Avic/EGFP=pballE}2.1* (gift from A. Herzig) was crossed with *P{Sgs3-GAL4.PD}* (BN 6870) and third instar larvae from the progeny were used. For extra sex combs analysis, mutant *ball^2^* and *w^1118^* were crossed to *Pc* alleles (*Pc^1^* and *Pc^XL5^*) at 25°C. Males in the progeny of these crosses were scored for extra sex comb analysis as described previously (Tariq et al., 2009). *ball^2^* and *w^1118^* were also crossed to *trx* alleles (*trx^1^* and *trx^E2^*) and with each other as control at 29°C. The progeny of these crosses were scored for *trx* mutant phenotypes. For analyzing expression of homeotic genes in vivo, embryos homozygous for *ball^2^* and *w^1118^*were used to stain with antibodies as described previously (Umer et al., 2019).

### *Drosophila* cell culture

Schneider’s *Drosophila* medium (Gibco, ThermoFisher Scientific), supplemented with 10% fetal bovine serum (Gibco, ThermoFisher Scientific) and 1% penicillin–streptomycin (Gibco, ThermoFisher Scientific) was used to culture *Drosophila* S2 cells. Express Five SFM (Gibco, ThermoFisher Scientific) supplemented with 20mM GlutaMAX (Gibco, ThermoFisher Scientific) and 1% penicillin–streptomycin was used to grow D.Mel-2 cells.

### Generation of stable cells, RNA isolation, Chromatin immunoprecipitation

To generate stable cell line with inducible expression vector of FLAG tagged BALL, *w^1118^*embryos were used to prepare cDNA and *ball* CDS was amplified. The *ball* CDS was first cloned in pENTR-d-TOPO™ entry vector (Thermo Fisher Scientific) followed by cloning in *pMTWHF* destination vector from DGVC (*Drosophila* Gateway Vector Collection) by setting up LR Clonase reaction following manufacturer’s protocol (ThermoFisher Scientific). The resulting *pMT-ball-FLAG* vector was transfected into S2 cells using Effectene transfection reagent (Qiagen) and stable cell line was generated by following the manufacturer’s instructions. The procedures employed for RNA isolation followed by expression analysis through qPCR as well as for ChIP and Western blotting using *Drosophila* cells are described previously (Umer et al., 2019). Genomic localization was determined by high throughput sequencing of libraries generated with BGISEQ-500. The data was filtered and the adaptor sequences were trimmed using BGI software (Drmanac et al., 2010; Huang et al., 2017). The filtered data was mapped to *Drosophila* genome (dm3 assembly) using bowtie tool of Galaxy version (Langmead et al., 2009). To obtain input normalized profiles, the callpeak function of MACS version 2 of Galaxy was used (Zhang et al., 2008). LogM-value profiles were obtained from GEO for H3K4me3 (GSE20787), H3K27ac (GSE20779), PC (GSE104059) and TRX (GSM2175519).

### Immunostaining

Larvae expressing GFP tagged BALL in salivary glands were dissected to isolate the salivary glands. Polytene squashes were prepared and immunostainings were performed using standard protocol (Sullivan et al., 2000). Stage 15 embryos were dechorionated and GFP-negative embryos were selected for immunostaining using standard protocol (Sullivan et al., 2000).

### Microscopy

For observing extra sex comb phenotype, Olympus SZ51 stereomicroscope was used. For imaging loss of abdominal pigmentation phenotype, male flies of the desired genotype were transferred to 70% ethanol to dehydrate. The dehydrated flies were dissected under Olympus SZ51 stereomicroscope on a dissection slide. The abdominal portion of the fly was isolated and rehydrated in water for 5-10 minutes. After rehydration, abdomens were mounted on a glass slide in Hoyer’s medium. A coverslip was placed over the specimen and was incubated at 65°C for 50 minutes. The final imaging was done using Nikon C-DSS230 epifluorescent stereomicroscope at 3.5× magnification. The same epifluorescent stereomicroscope was used for the selection of GFP negative embryos (homozygous *ball^2^* mutant embryos) in experiments where immunostaining and real-time PCR analysis of embryos was performed. Imaging of embryos was done at 20× magnification on Nikon C2 confocal microscope. NIS elements image acquisition software was utilized for imaging and analysis. All larval dissections for polytene chromosomes were performed using LABOMED CZM6 stereo zoom microscope and Nikon C2 confocal microscope was used for imaging. Zooming to 60× was used for visualization of polytene chromosomes.

### Antibodies

Following antibodies were used during this study: mouse anti-Abd-B (DSHB, 1A2E9, IF: 1:40), rabbit anti-TRX (gift from R. Paro, IF: 1:20), rabbit anti-PC (Santa Cruz, D220, IF: 1:20), rabbit anti-BALL (Gift from A. Herzig, IF 1:20, ChIP: 2μl) mouse anti-GFP (Roche, 11814460001, IF:1:50), mouse anti-Tubulin (Abcam, ab44928, WB: 1:2000), mouse anti-FLAG M2 (Sigma Aldrich, WB: 1:2000, ChIP: 5μl), mouse anti-H2A-ubi (Millipore, 05-678, ChIP: 5μl). HRP conjugated secondary antibodies (Abcam) were used at 1:10,000 dilution for western blotting while cy3 and Alexa Fluor 488 conjugated secondary antibodies (Thermo Fisher Scientific) were used at 1:100 dilution.

## Supplementary Materials

Figure S1 Schematics and data normalization of kinome-wide screen.

Figure S2. ChIP peak coverage of BALL across genome.

Table S1. List of peaks generated using ChIPseeker and presented along with their coordinates.

---

**Figure S1.**
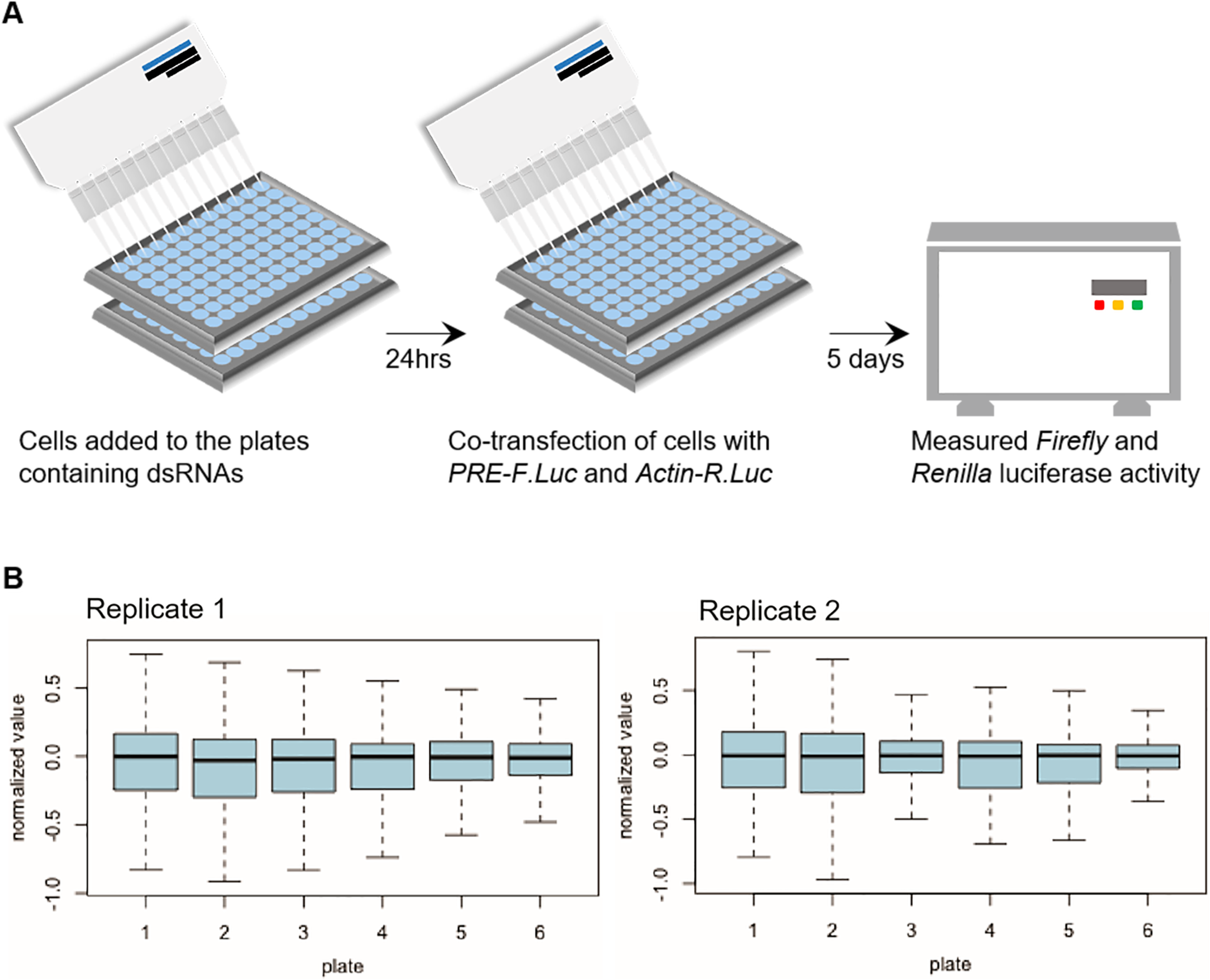
Schematics of ex vivo kinome-wide RNAi screen and data analysis. (A) 384-well plates containing dsRNAs against different genes were loaded with equal number of cells in each well. 24 hours after seeding cells, *PRE-F.Luc* and *Actin-R.Luc* were co-transfected and luciferase values were determined five days later. (B) Box plots for plate median normalized data for kinome-wide RNAi screen, replicate 1 (left) and replicate 2 (right).

**Figure S2.**
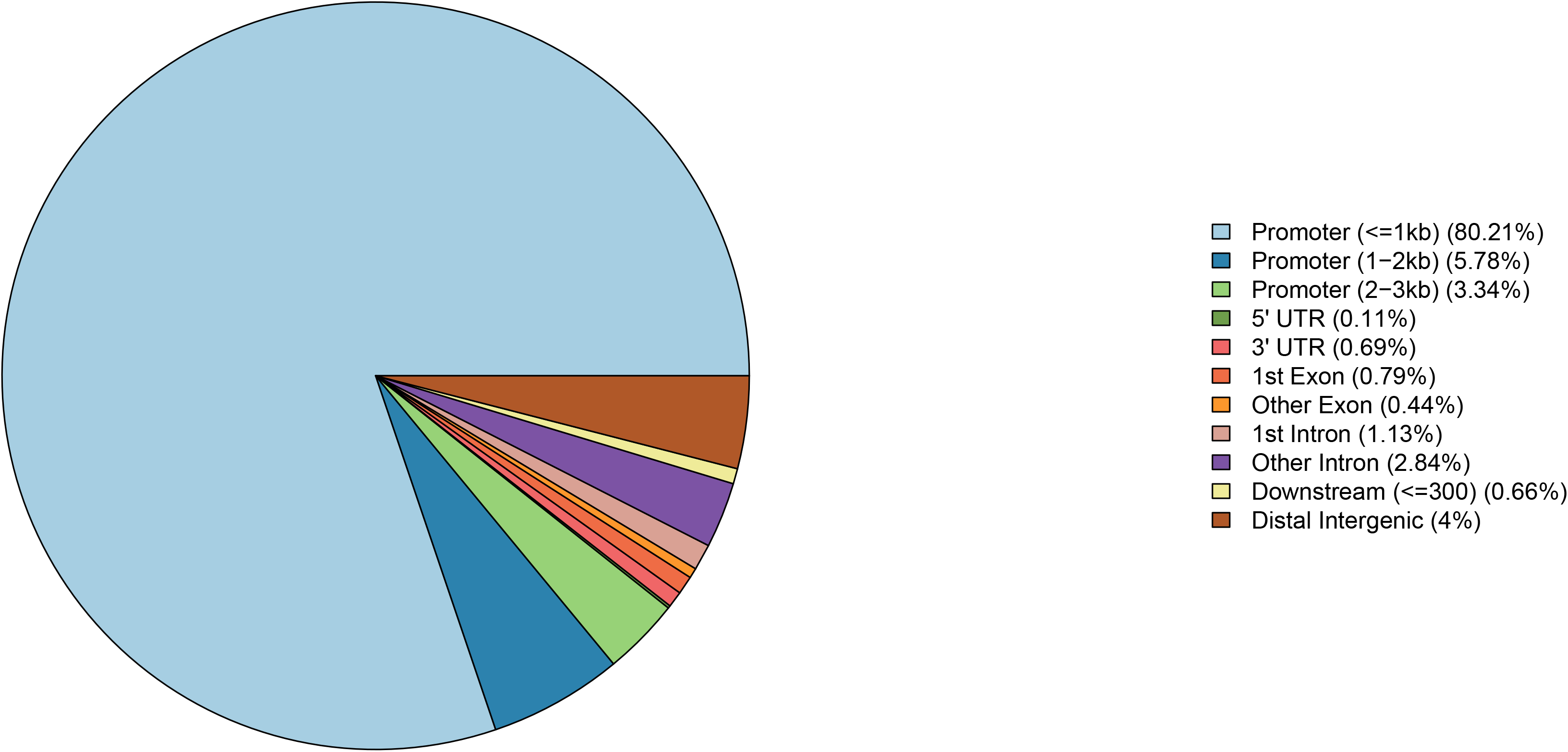
ChIP peak coverage of BALL across genome. ChIPseeker is utilized to generate a pie chart that represent distribution of BALL across different regions of genome. BALL is enriched predominantly in promoter (<=1kb) regions followed by promoter (1−2kb) regions, distal intergenic regions and promoter (2−3kb) regions (Yu et al., 2015).

**Table S1. List of peaks generated using ChIPseeker and presented along with their coordinates**.

## Declarations

## Acknowledgements

We would like to thank Michael Boutros at DKFZ, Germany for providing access to all the facilities for RNAi screen and Alf Herzig for providing us Ballchen flies and anti-BALL antibodies.

## Author contributions

MHFK, JA, ZU, NS, SA and MT designed research and MHFK, JA, ZU, NS, AS and SA performed experiments. MHFK, JA, ZU, NS and MT wrote the manuscript. AM analyzed the screen data. All authors approved the final version of the manuscript.

## Data and materials availability

All data generated and analyzed in the study are available in the main or additional files provided.

---

**Table S1.**
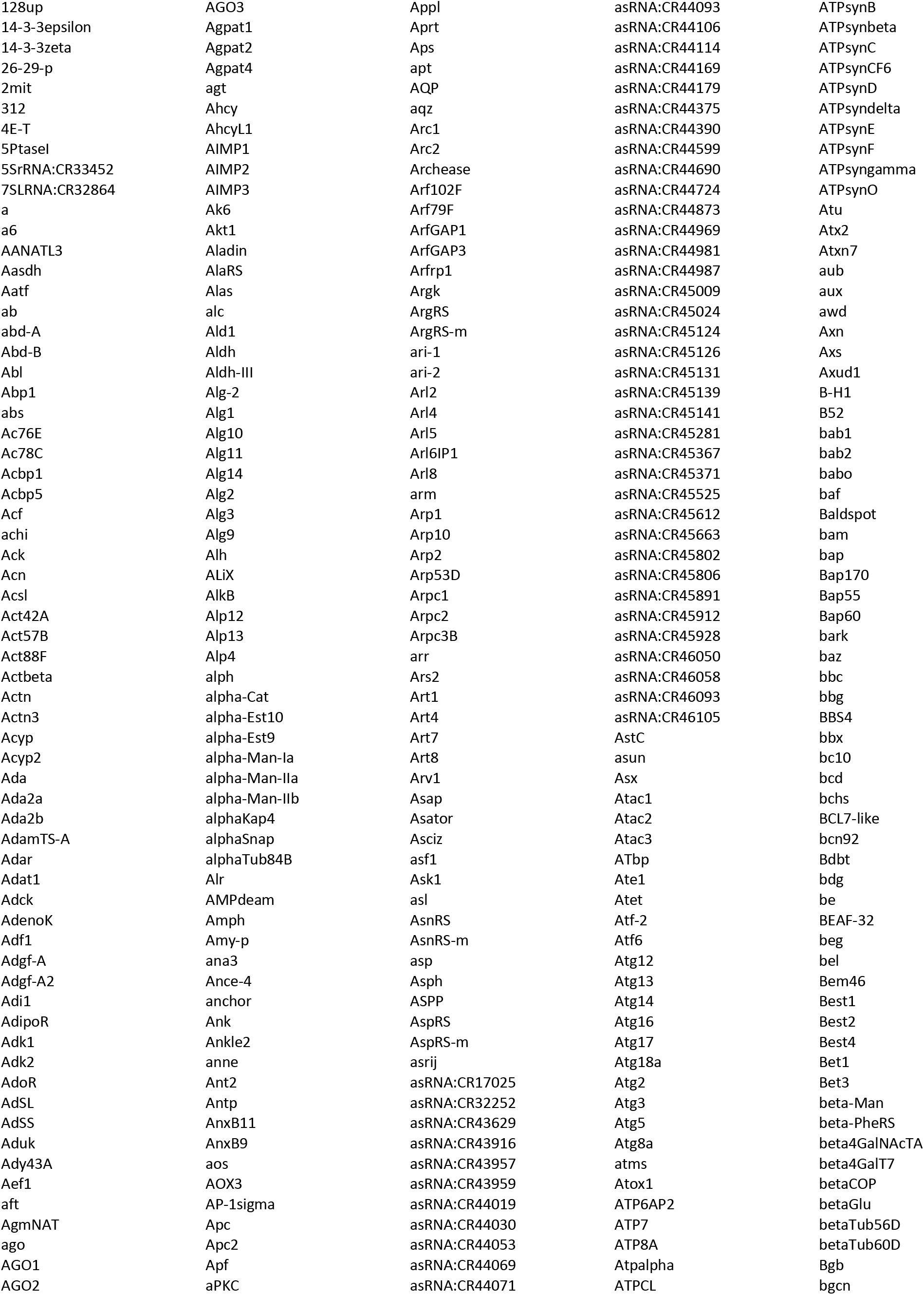
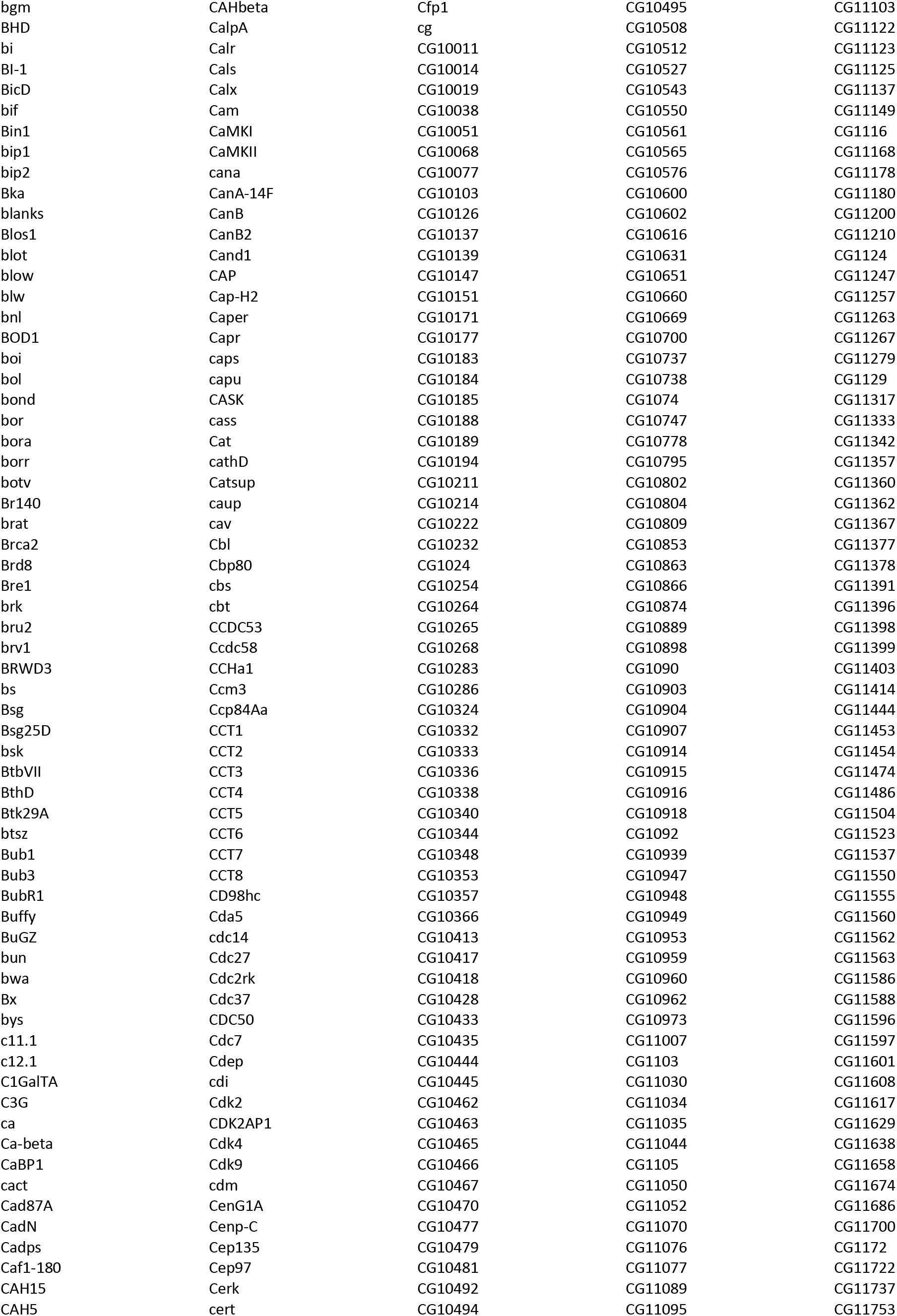
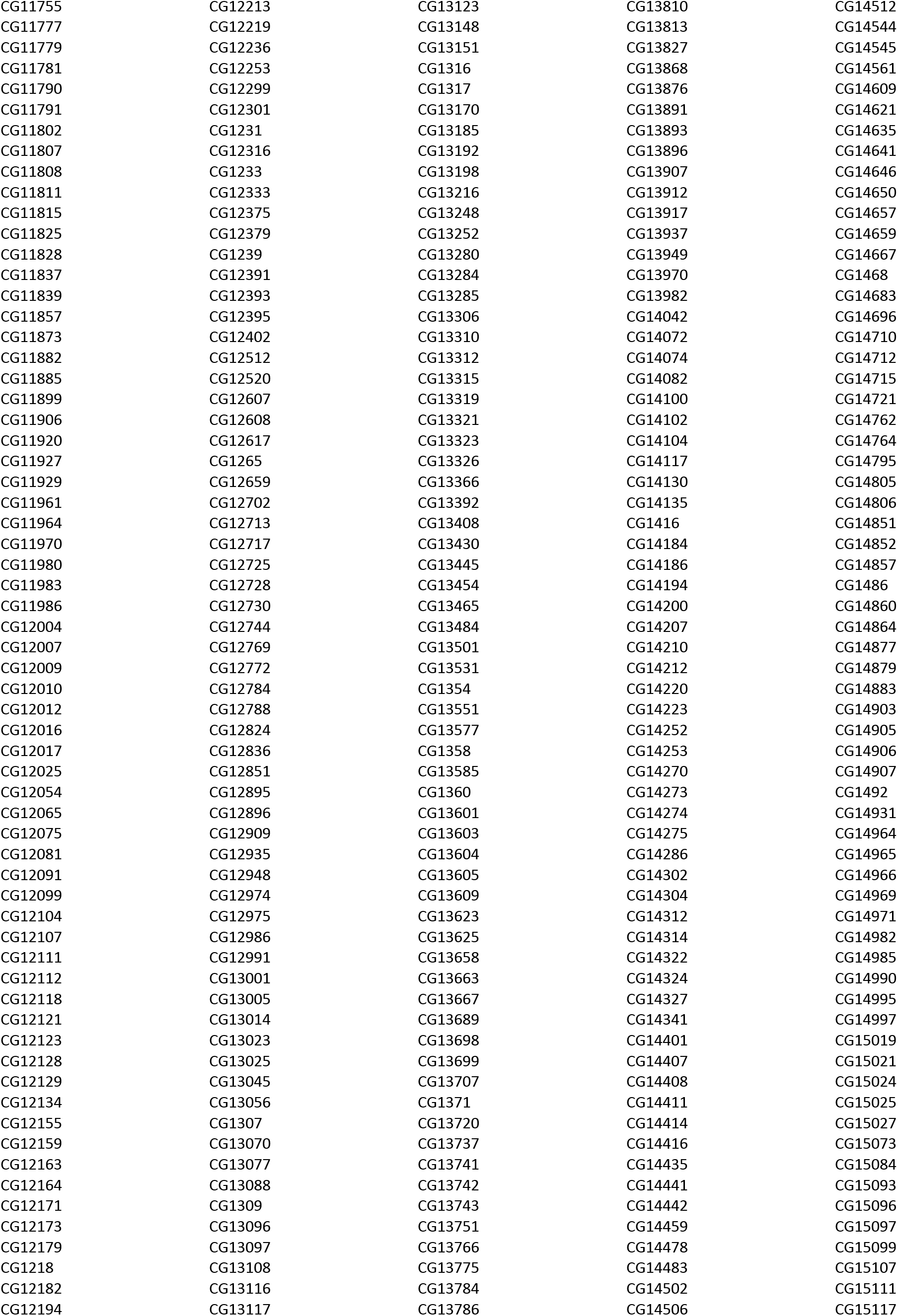
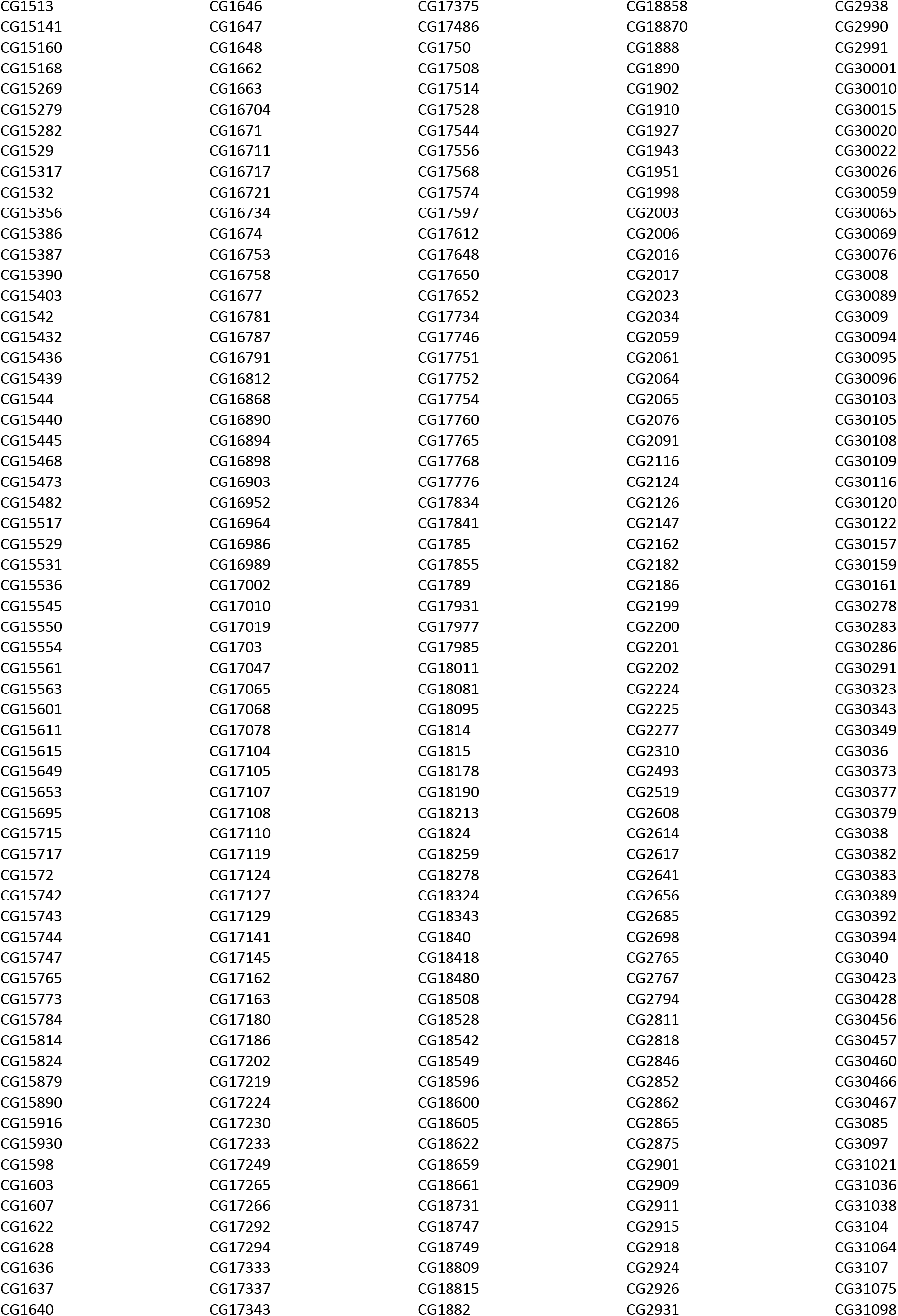
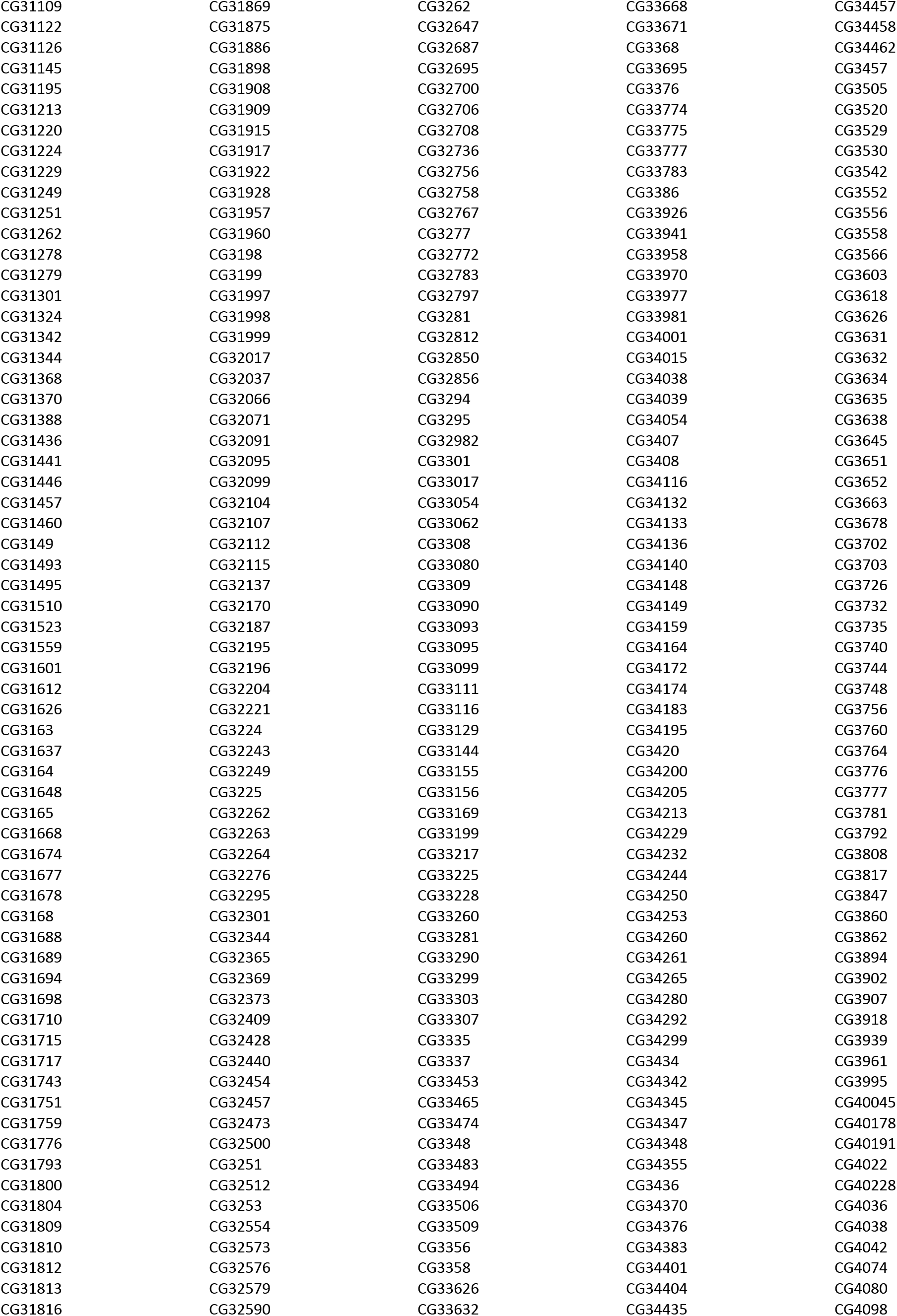
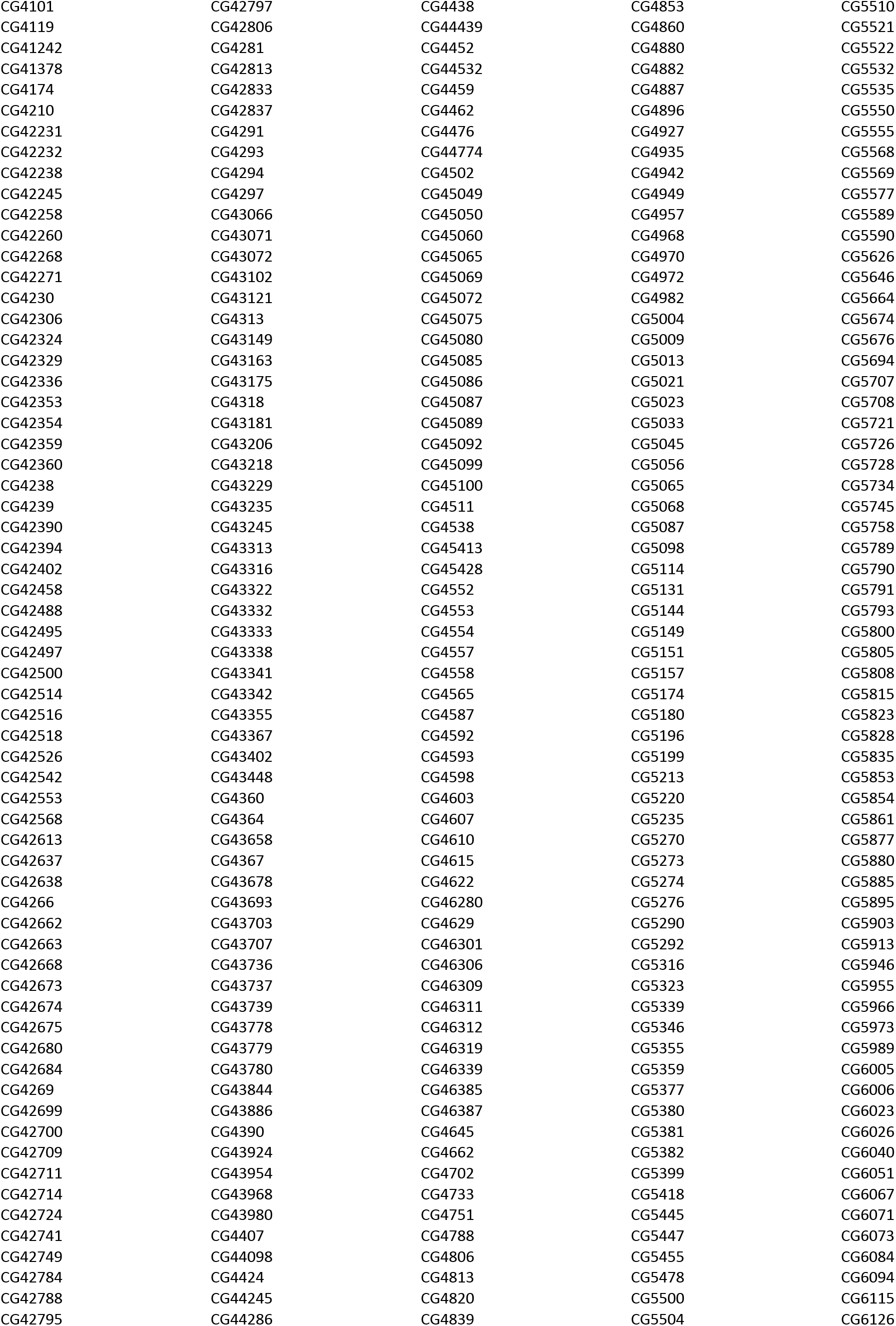
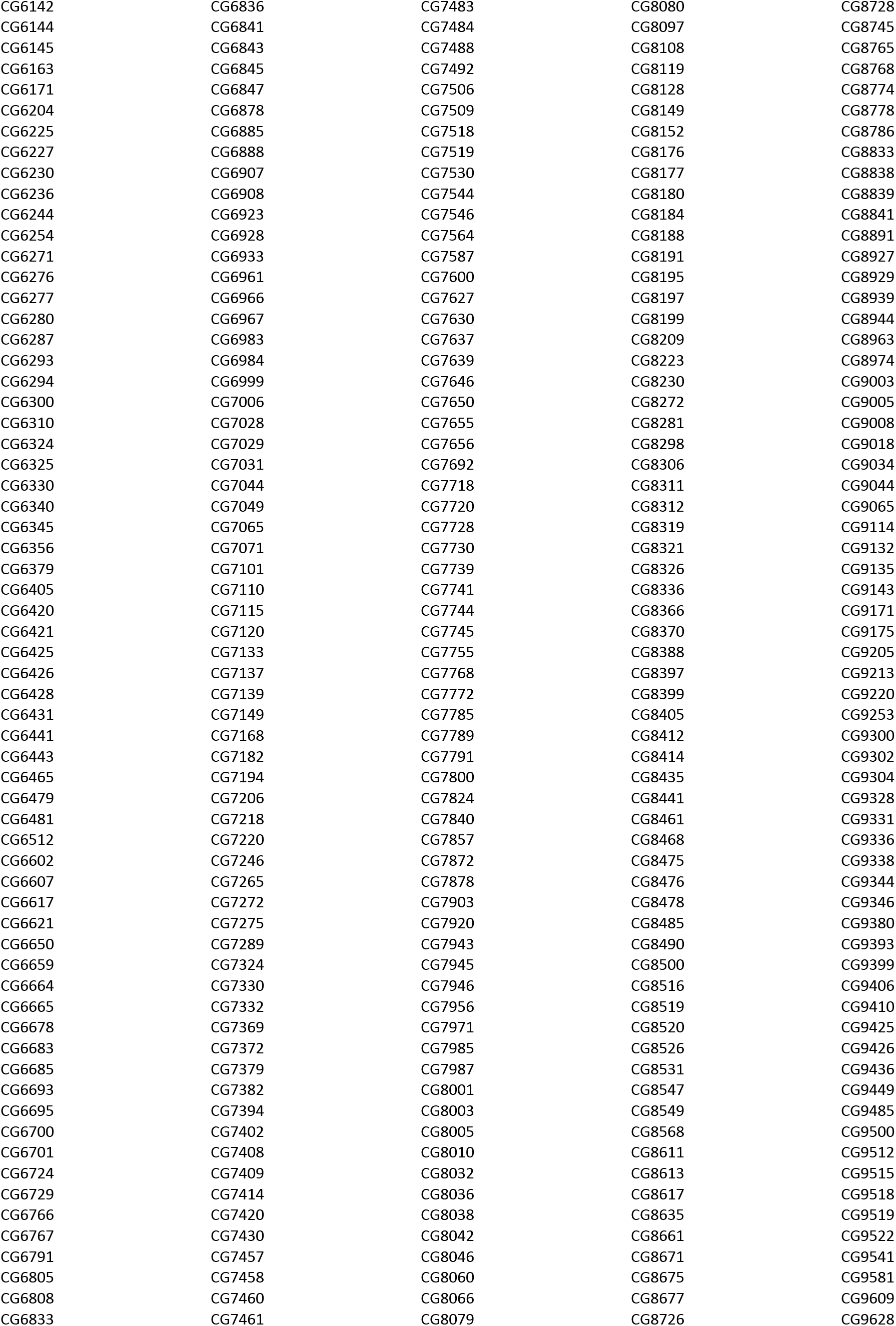
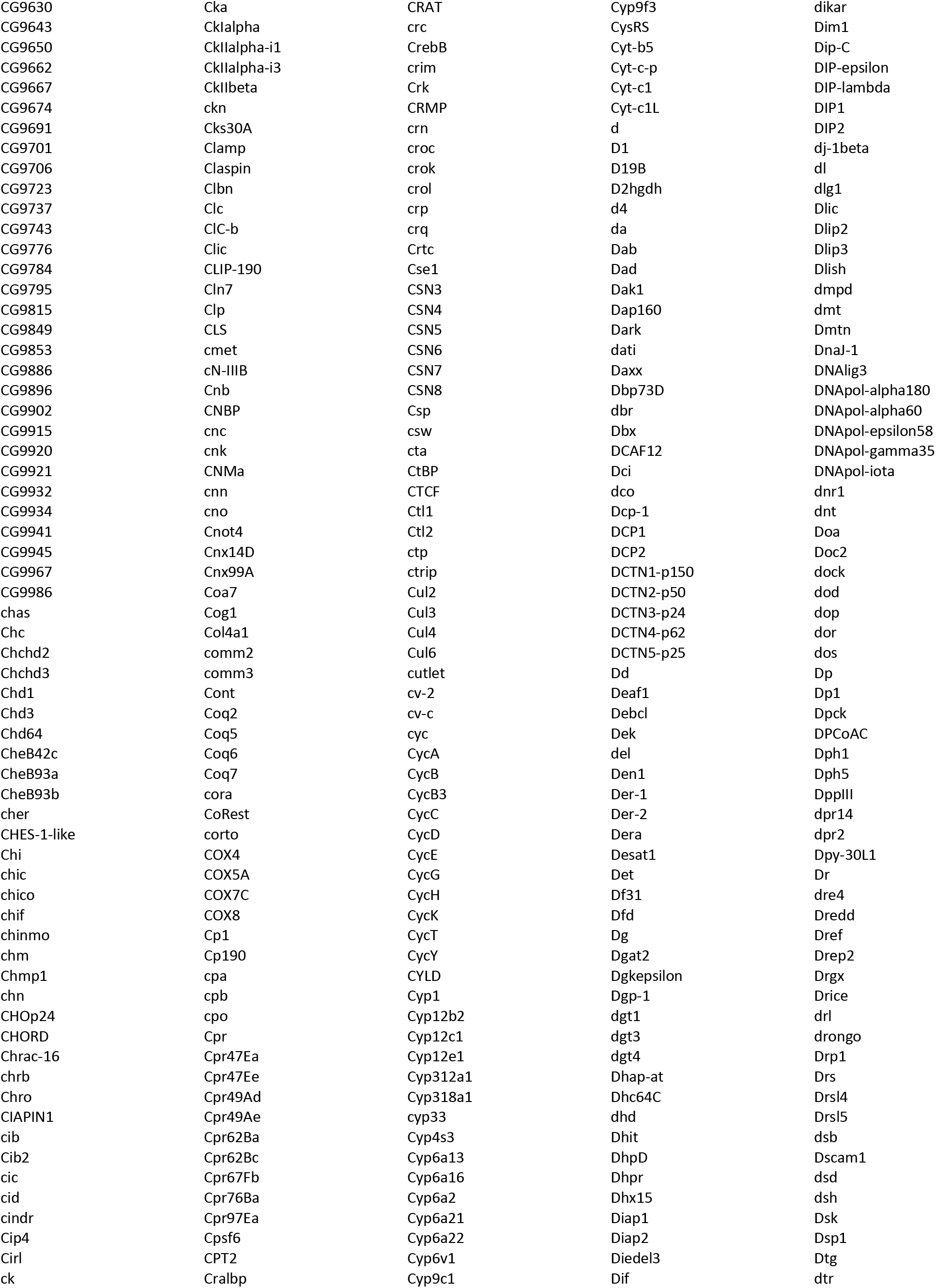
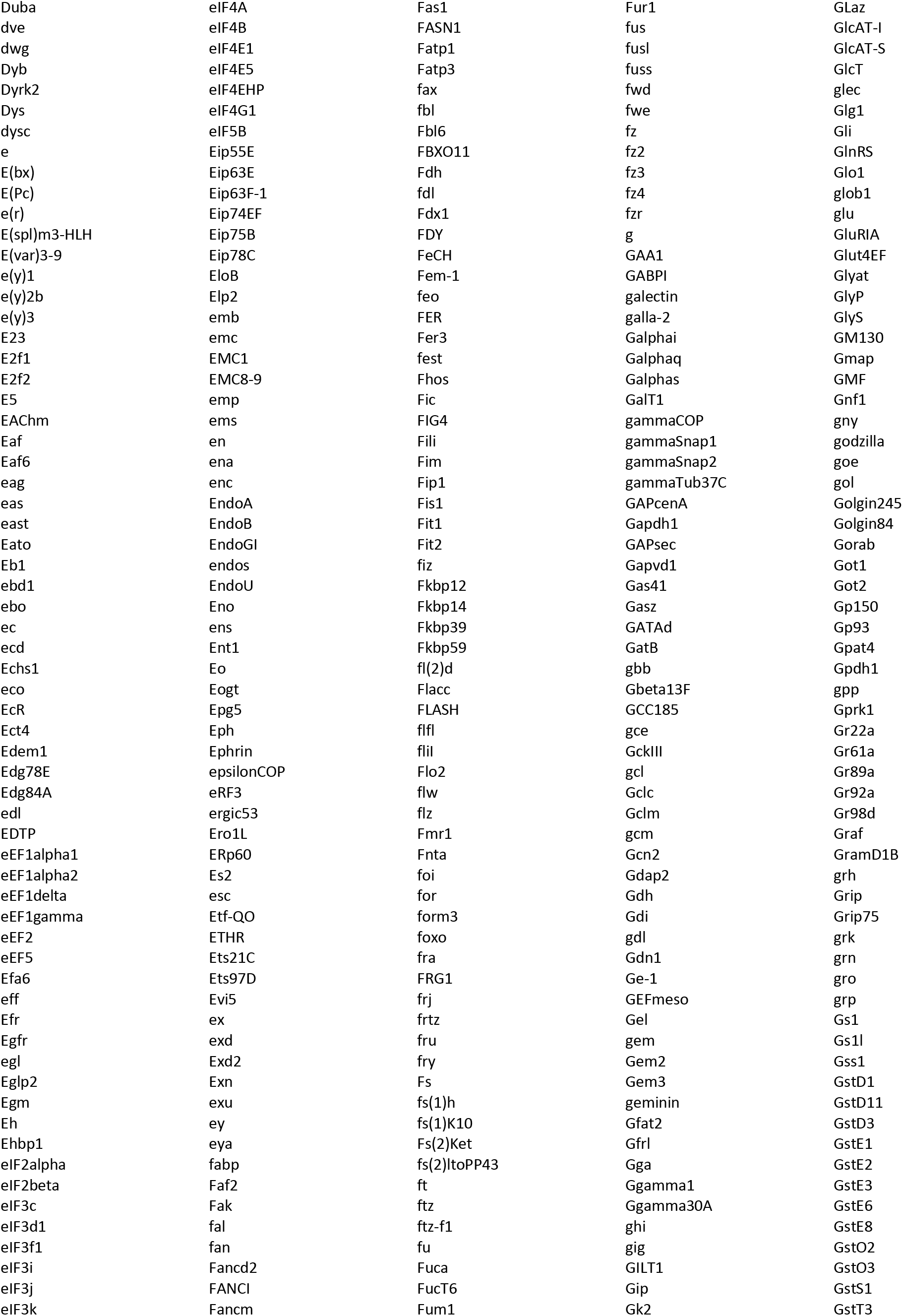
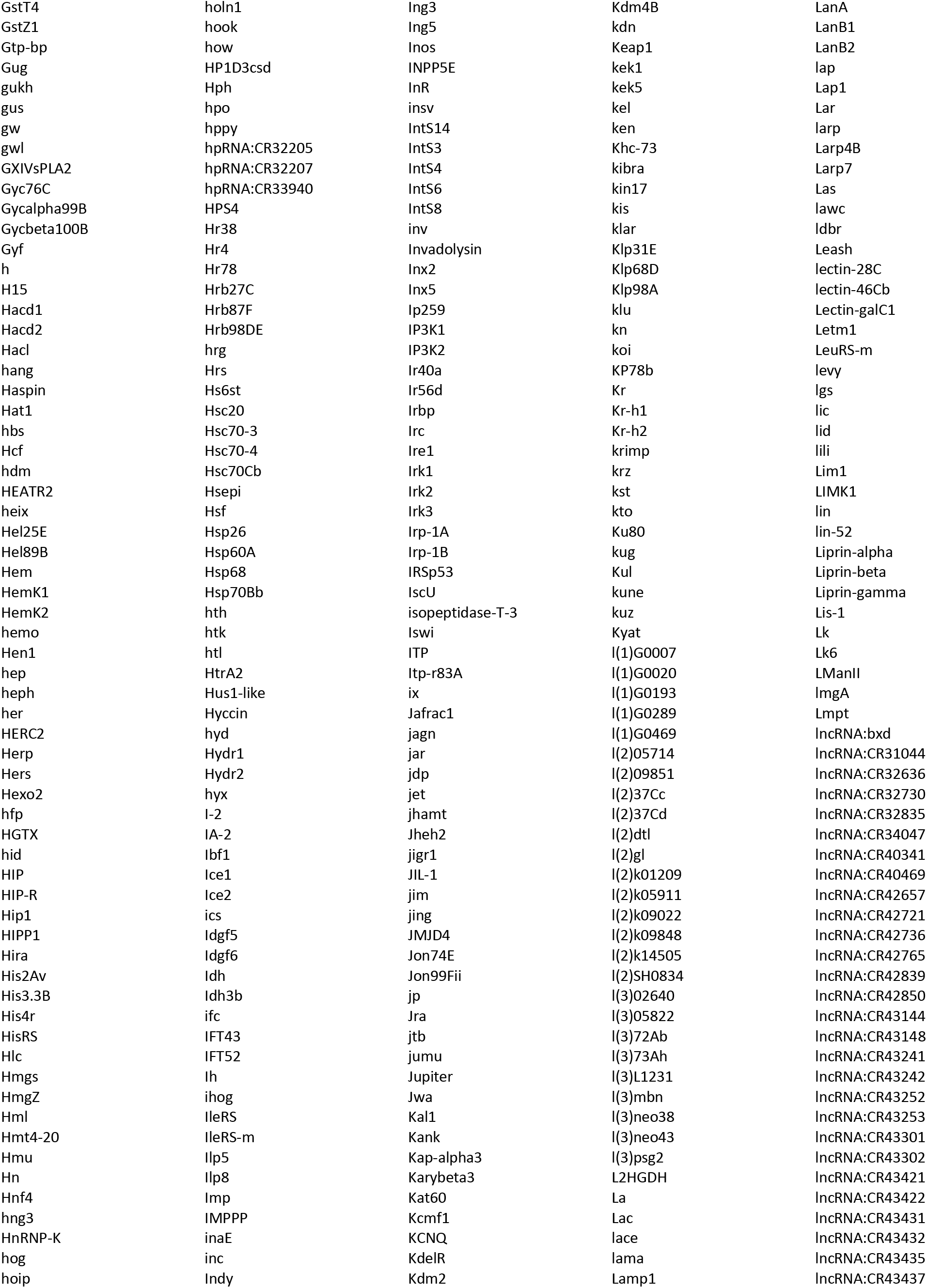
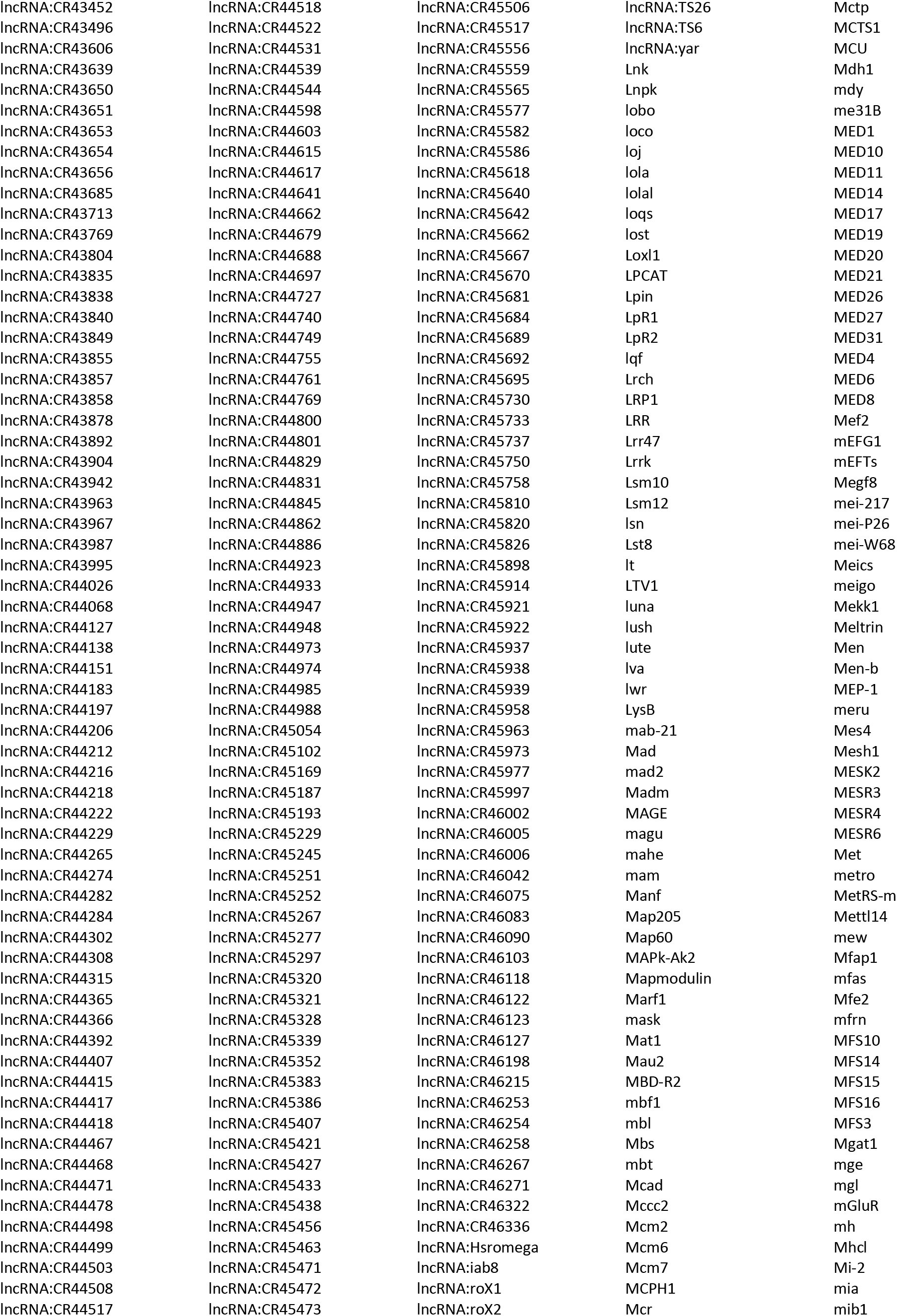
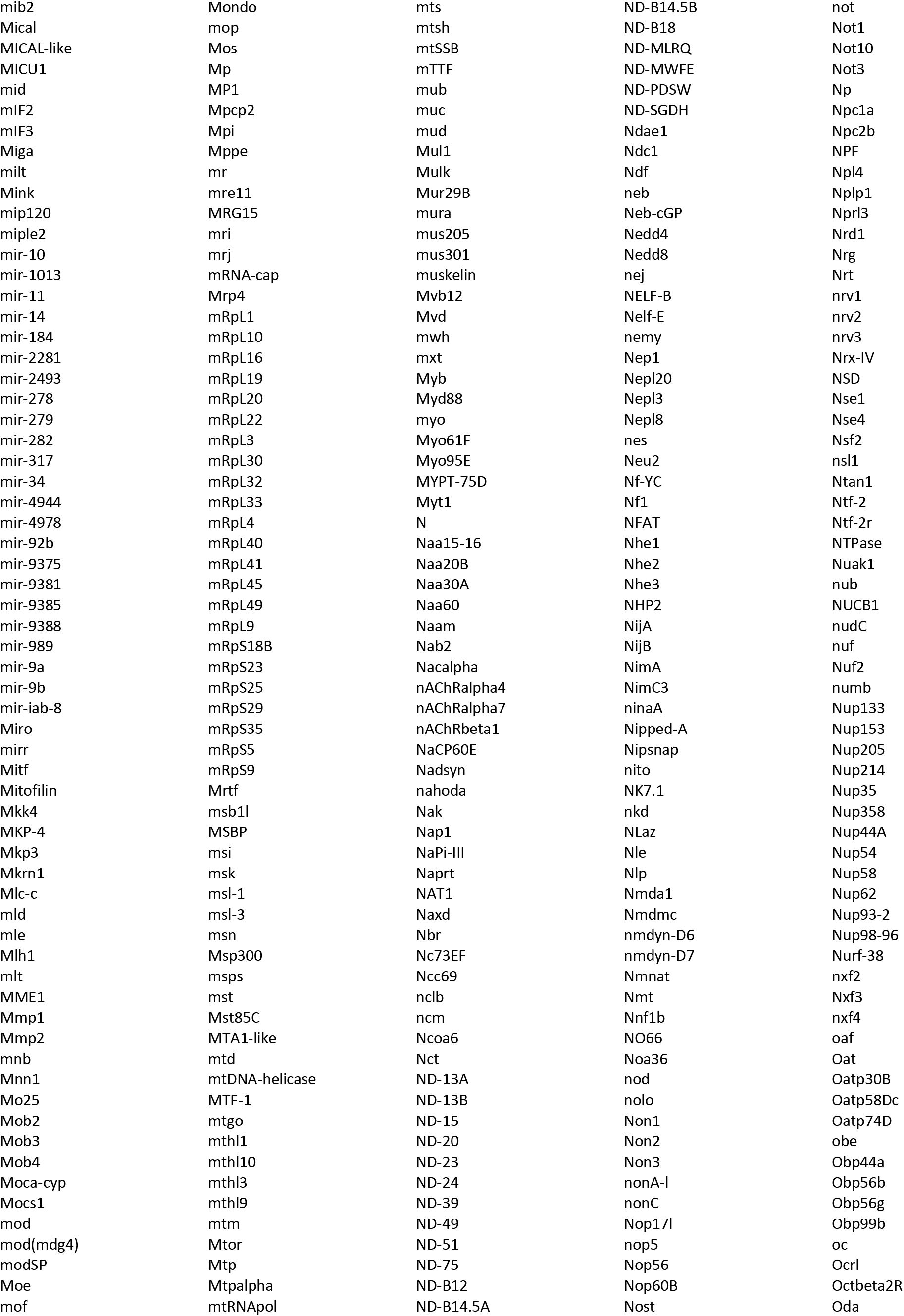
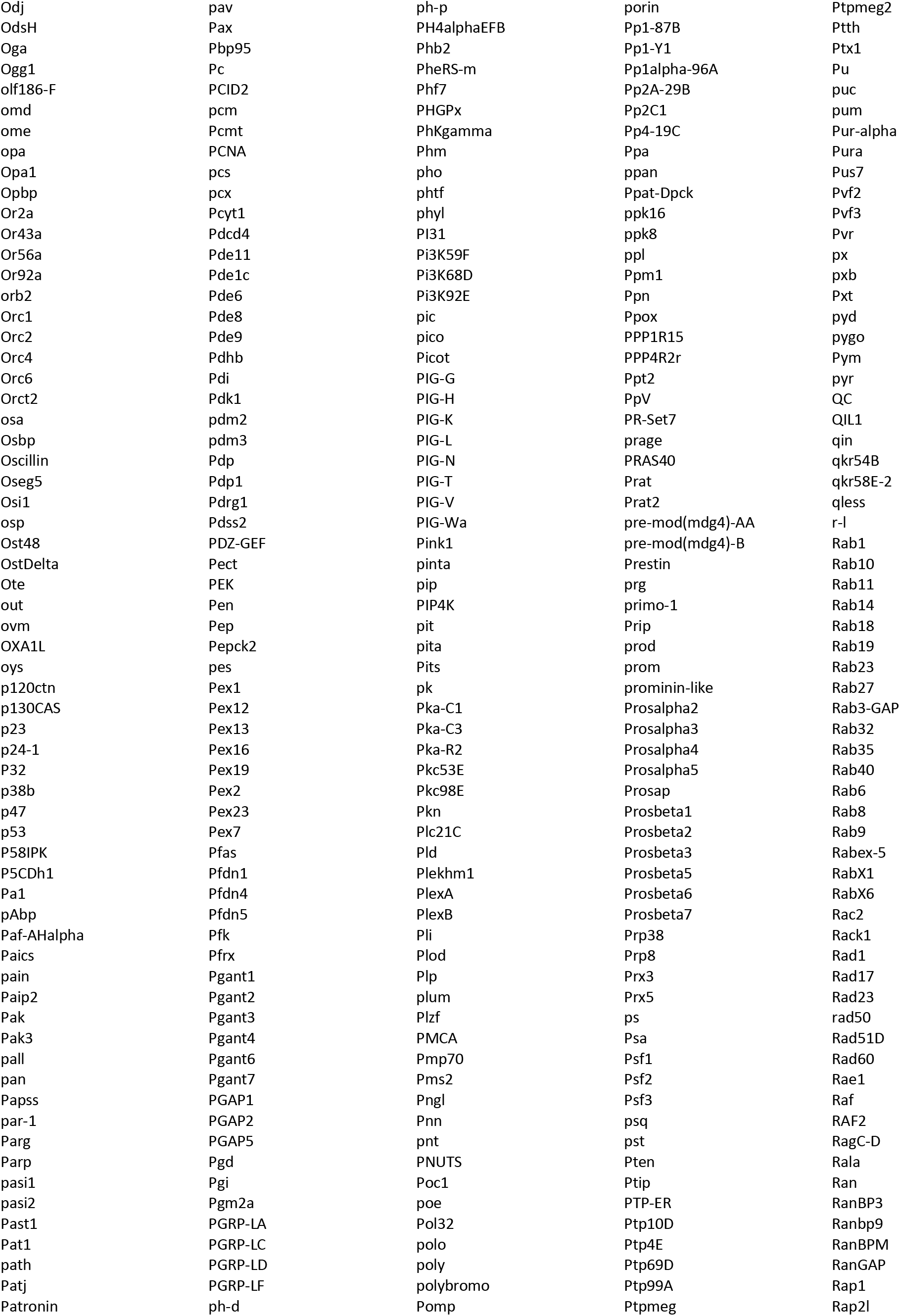
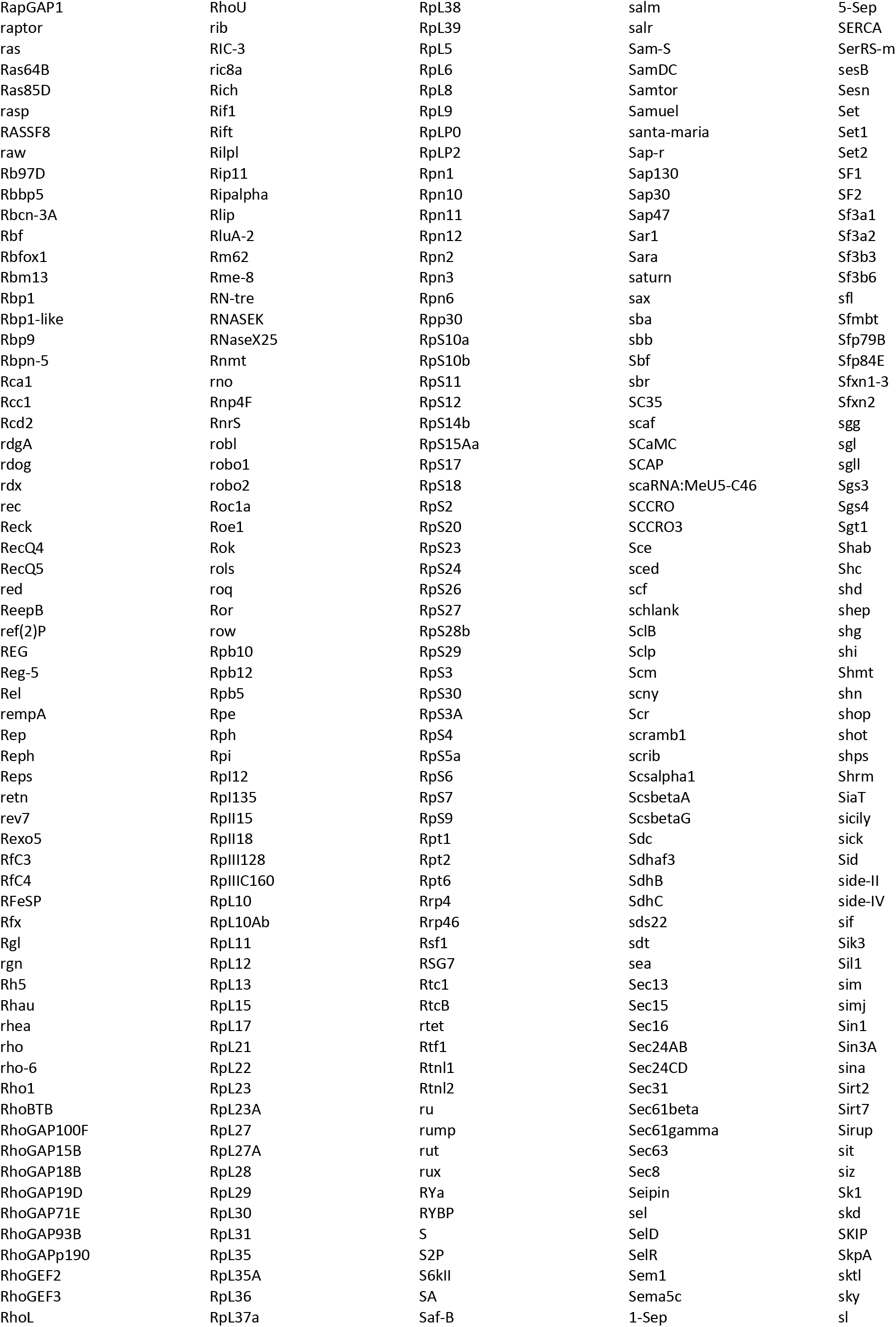
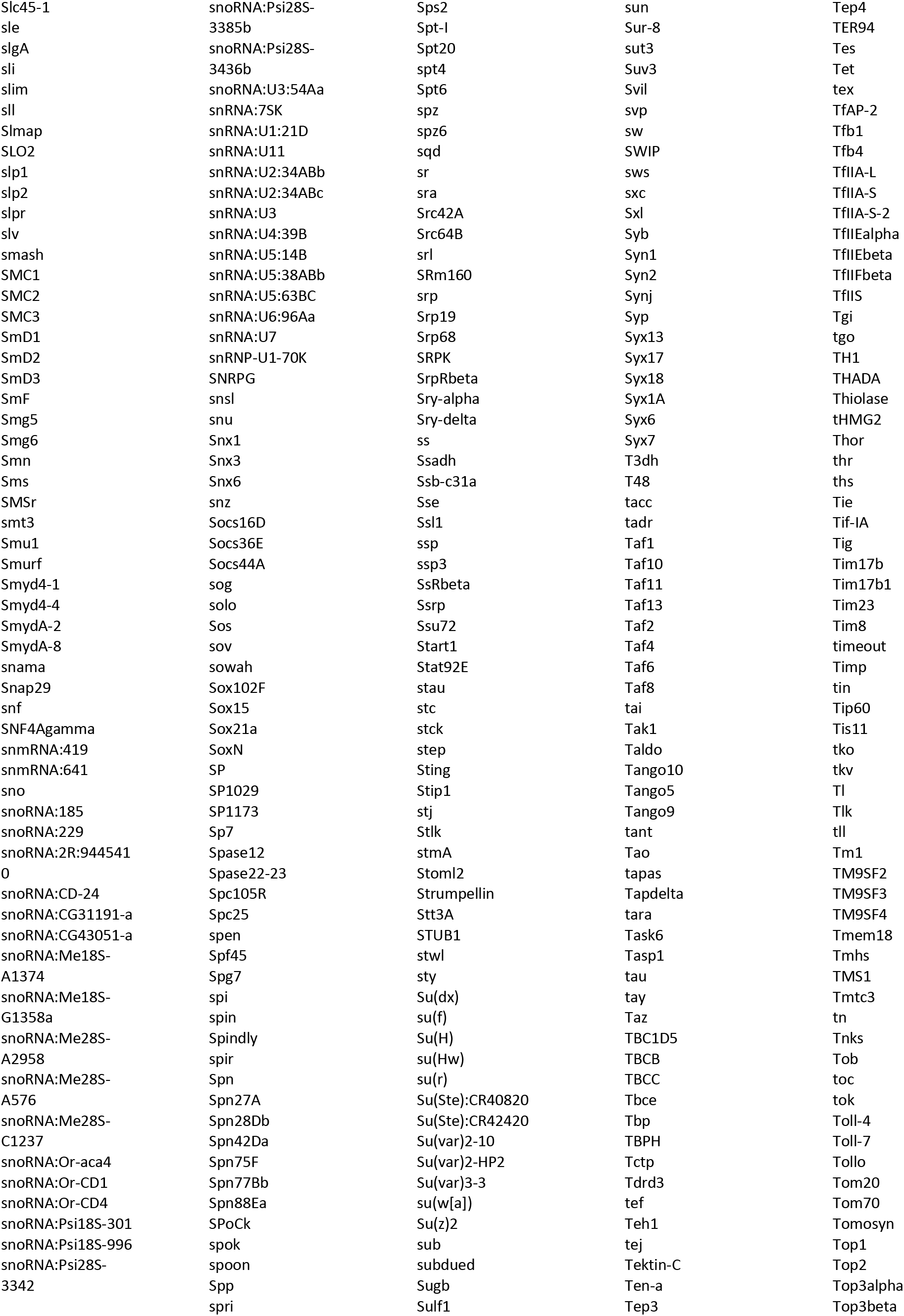
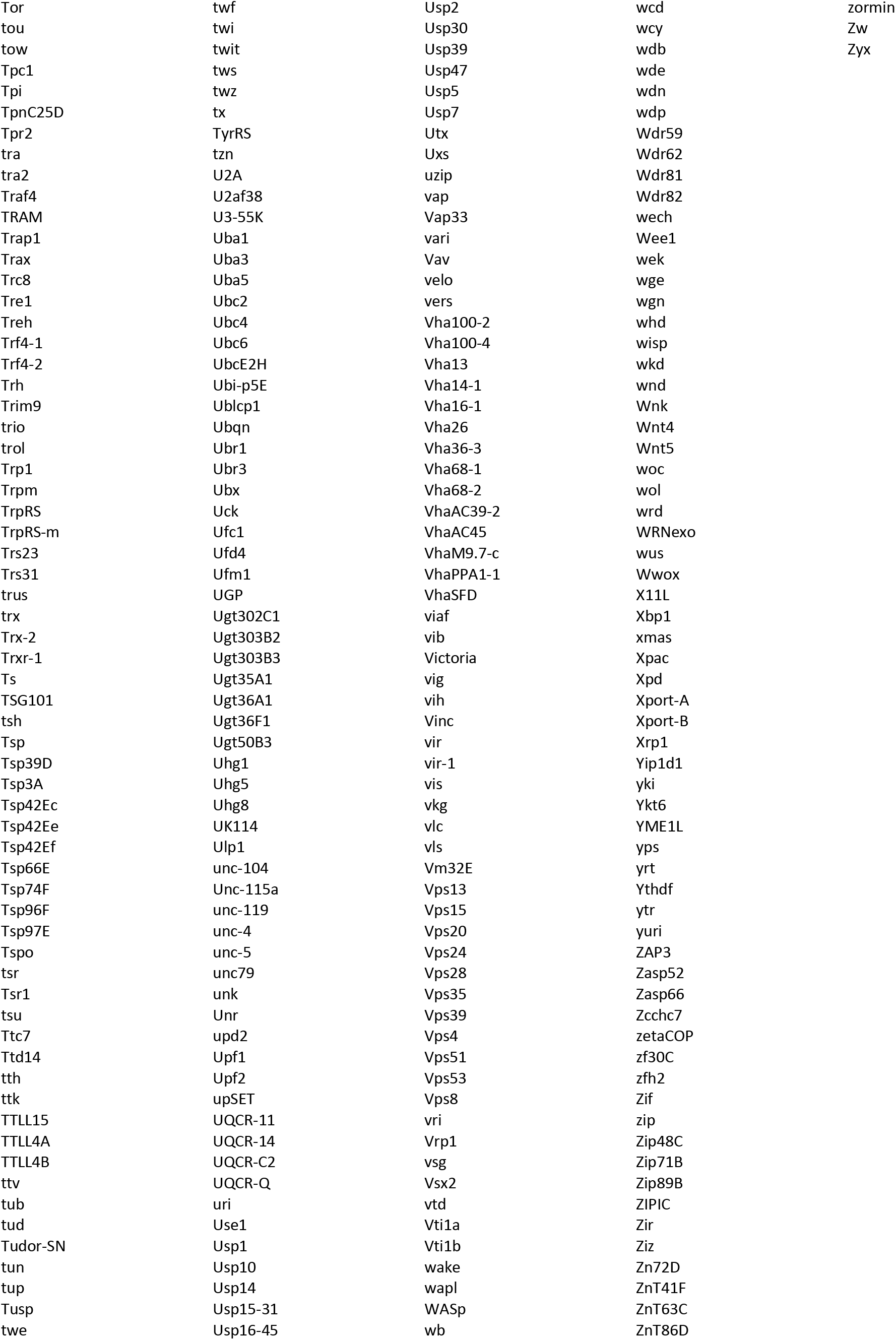
List of genes where BALL peaks are present in genome-wide binding profile. Gene names or annotation numbers are given.

